# Low-N protein engineering with data-efficient deep learning

**DOI:** 10.1101/2020.01.23.917682

**Authors:** Surojit Biswas, Grigory Khimulya, Ethan C. Alley, Kevin M. Esvelt, George M. Church

**Affiliations:** Wyss Institute for Biologically Inspired Engineering, Harvard University; Nabla Bio, Inc.; Telis Bioscience Inc.; MIT Media Lab, Massachusetts Institute of Technology; Department of Genetics, Harvard Medical School

## Abstract

Protein engineering has enormous academic and industrial potential. However, it is limited by the lack of experimental assays that are consistent with the design goal and sufficiently high-throughput to find rare, enhanced variants. Here we introduce a machine learning-guided paradigm that can use as few as 24 functionally assayed mutant sequences to build an accurate virtual fitness landscape and screen ten million sequences via *in silico* directed evolution. As demonstrated in two highly dissimilar proteins, avGFP and TEM-1 β-lactamase, top candidates from a single round are diverse and as active as engineered mutants obtained from previous multi-year, high-throughput efforts. Because it distills information from both global and local sequence landscapes, our model approximates protein function even before receiving experimental data, and generalizes from only single mutations to propose high-functioning epistatically non-trivial designs. With reproducible >500% improvements in activity from a single assay in a 96-well plate, we demonstrate the strongest generalization observed in machine-learning guided protein function optimization to date. Taken together, our approach enables efficient use of resource intensive high-fidelity assays without sacrificing throughput, and helps to accelerate engineered proteins into the fermenter, field, and clinic.

## Introduction

Protein engineering holds great promise for nanotechnology, agriculture, and medicine. However, design is limited by our ability to search through the vastness of protein sequence space, which is only sparsely functional^1,2^. When searching for high functioning sequences, engineers must be wary of the pervasive maxim, “you get what you screen for”, which cautions against over-optimizing a protein’s sequence using functional assays that may not be fully aligned with the final design objective^3–6^. However, in most resource-constrained real-world settings, including the design of protein therapeutics^7,8^, agricultural proteins^9^, and industrial biocatalysts^10,11^, engineers must often compromise assay fidelity (careful endpoint-resembling measurements of a small number of variants) for assay throughput (high-throughput proxy measurements for a large number of variants)^12,13^. Consequently, the best candidates identified by early stage high-throughput (>10^4^ variants) proxy experiments^9,11,14^ will often fail in validation under higher-fidelity, later stage assays^13,15–17^. Moreover, high-throughput assays do not exist at all for many classes of proteins, making them inaccessible to screening and directed evolution^18–24^.

Here we focus on enabling large-scale exploration of sequence space using only a small number — “low-N” — of functionally characterized training variants. We recently developed UniRep^25^, a deep learning model trained on a large unlabeled protein sequence dataset. From scratch and from sequence alone, UniRep learned to distill the fundamental features of a protein — including biophysical, structural, and evolutionary information — into a holistic statistical summary, or *representation*.

We reasoned that combining UniRep’s global knowledge of functional proteins with just a few dozen functionally characterized mutants of the target protein might suffice to build a high-quality model of a protein’s fitness landscape. Combined with *in silico* directed evolution, we hypothesized that we could computationally explore these landscapes at a scale of 10^7^-10^8^ variants, rivalling even the highest-throughput screens. Here, we test this paradigm in two fundamentally different proteins — a eukaryotic green fluorescent protein from *Aequorea victoria* (avGFP), and a prokaryotic β-lactam hydrolyzing enzyme from *Escherichia coli* (TEM-1 β-lactamase). We demonstrate reliable production of substantially optimized designs with just 24 or 96 characterized sequence variants as training data.

## Results

### A paradigm for low-N protein engineering

To meet the enormous data requirement of supervised deep learning — typically greater than 10^6^ labeled data points^26,27^— current machine learning-guided protein design approaches must gather high-throughput experimental data^28–31^ or abandon deep learning altogether^18,20,21,32–37^. We reasoned that UniRep could leverage its existing knowledge of functional protein sequences to substantially reduce this prohibitive data requirement and enable low-N design.

For low-N engineering of a given target protein, our approach features five steps (Fig. 1):

1. Global unsupervised pre-training of UniRep on >20 million raw amino acid sequences to distill general features of all functional proteins, as described previously^25^ (Fig. 1a).
2. Unsupervised fine-tuning of UniRep on sequences evolutionarily related to the target protein (evotuning) to learn the distinct features of the target family. We call this model, which combines features from both the global and local sequence landscape, evotuned UniRep, or eUniRep (Fig. 1b).
3. Functional characterization of a low-N number of random mutants of the wild-type target protein to train a simple supervised top model that uses eUniRep’s representation as input (Fig. 1c). Together, eUnirep and the top model define an end-to-end sequence-to-function model that serves as a surrogate of the protein’s fitness landscape.
4. Markov Chain Monte Carlo-based *in silico* directed evolution on this surrogate landscape (Fig. 1d-e).
5. Experimental characterization of top sequence candidates that are predicted to have improved function relative to wild-type (>WT).

**Figure 1.**
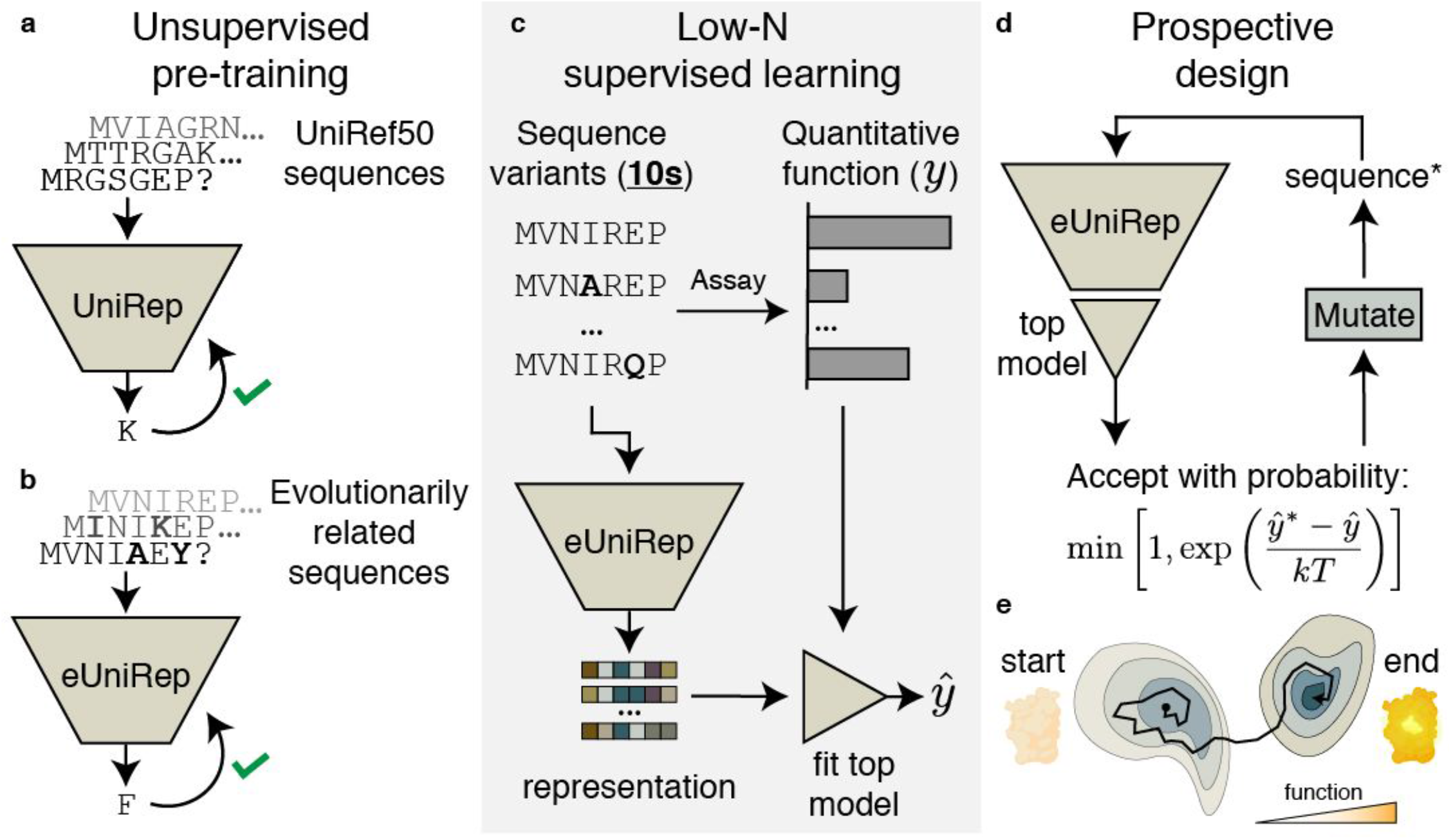
UniRep-guided *in silico* directed evolution for low-N protein engineering. **a)** UniRep is globally trained on a large sequence database (UniRef50) as described previously^25^. **b)** This trained, unsupervised model is further fine-tuned to sequences that are evolutionarily related to the protein of engineering interest (eUniRep). **c)** A low-N number of mutants are obtained, characterized, and used to train regularized linear regression “on top” of eUniRep’s representation. **d)** *In silico* directed evolution is used to navigate this virtual fitness landscape and propose putatively optimized designs that are then experimentally characterized. This design loop may be repeated until desired functionality is reached. **e)** Illustration of the evolutionary process.

To understand the utility of eUniRep’s global + local representation, we considered a control model which was trained *de novo* solely on the local sequence neighborhood^38–41^ of the target protein (Local UniRep). Thus, Local UniRep lacks global information about all known sequence space. As an additional control, we included one-hot encoding, as an explicit and exact flattened binary matrix representation of the full amino acid sequence (Full AA), to contextualize the importance of any evolutionary information (Methods).

We first evaluated our approach in retrospective experiments using pre-existing and newly designed datasets of characterized mutant proteins (Methods, Supplementary Fig. 1). We found that only globally pre-trained eUniRep enabled consistent low-N retrospective performance, and that with the right regularized top model, meaningful generalization required only 24 training mutants (Supplementary Fig. 2). Random selection of these 24 mutants from the output of error-prone PCR or single-mutation deep mutational scans worked as well as more tailored approaches (Methods).

**Figure 2.**
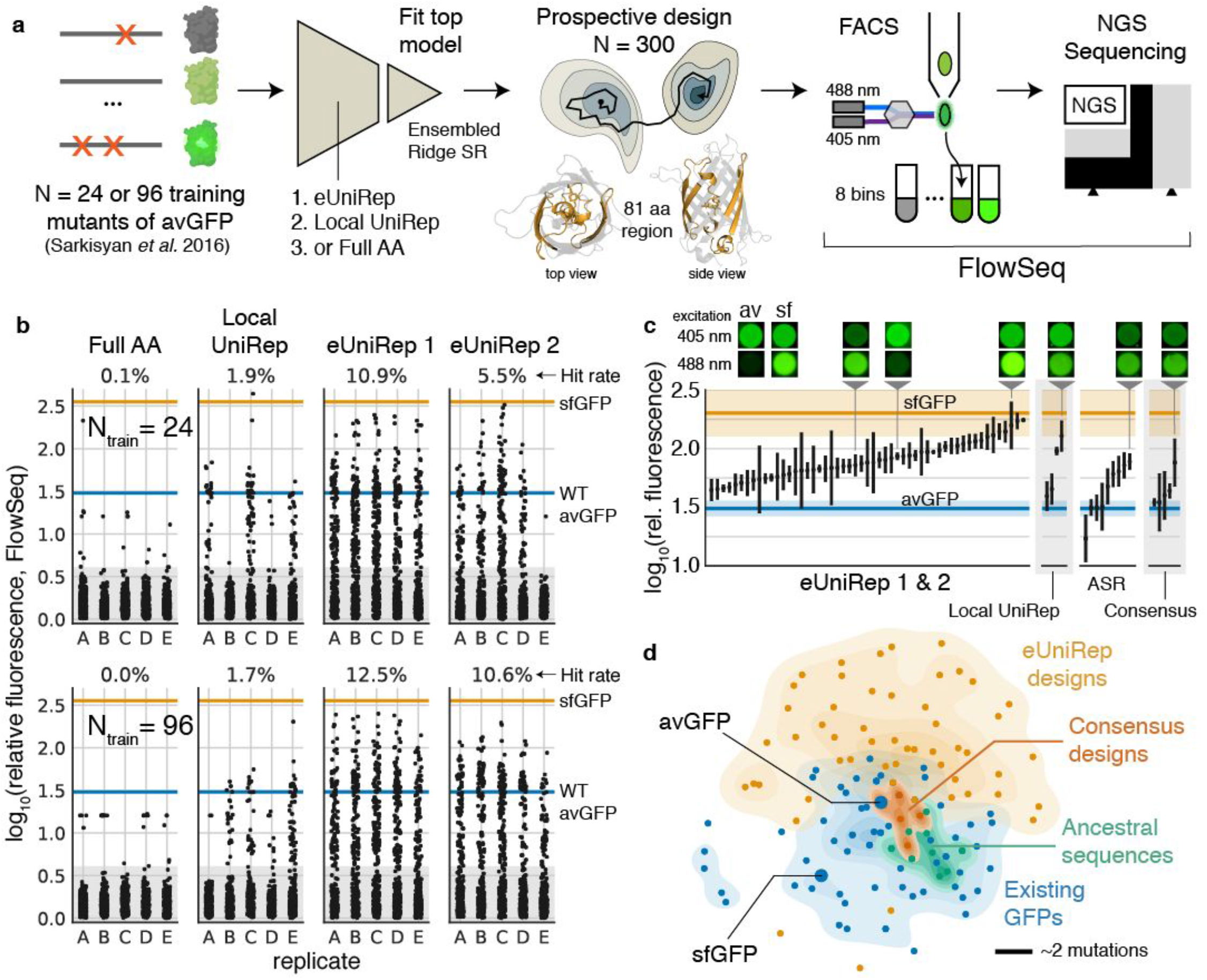
eUniRep enables low-N engineering of avGFP. **a)** Experimental workflow describing training mutant acquisition, sequence-to-function modeling, *in silico* directed evolution, and the use of FlowSeq to quantitatively characterize designs in multiplex. **b)** Low-N engineering results for 24 (top) and 96 (bottom) training mutants. eUniRep 1 and 2 correspond to two replicate evotunings initialized from the same globally pre-trained UniRep. **c)** Quantitative flow-cytometric measurements of top eUniRep and Local UniRep designs, as well as ASR and consensus sequence designs. Shown above are false-colored images of *E. coli* expressing avGFP (av), sfGFP (sf), and a subset of the designs under 405 nm or 488 nm excitation, read with a 525/50 emission filter. **d)** Distance-preserving multidimensional scaling plot illustrating the diversity of eUniRep designs compared to existing GFPs, ASRs, and consensus sequence designs. Scale bar of 2 mutations shown.

### Low-N engineering of the fluorescent protein avGFP

To test our approach prospectively, we attempted low-N optimization of the fluorescence intensity of the original green fluorescent protein from *Aequorea victoria* (avGFP) (Fig. 2a). The design process consisted of randomly sampling N=24 or N=96 training mutants from error-prone PCR^42^, representing sequences, training a top model, and performing *in silico* directed evolution to produce 300 putatively optimized designs within a 15 mutation “trust radius” of wild-type (Methods). We replicated this process 5 times for each N and representation model, yielding a total of 12,000 sequence designs. The design window spanned an 81 amino acid region of avGFP that included the central chromophore-bearing helix and four straddling beta-sheets (Fig. 2a; Methods; Supplementary Fig. 3).

**Figure 3.**
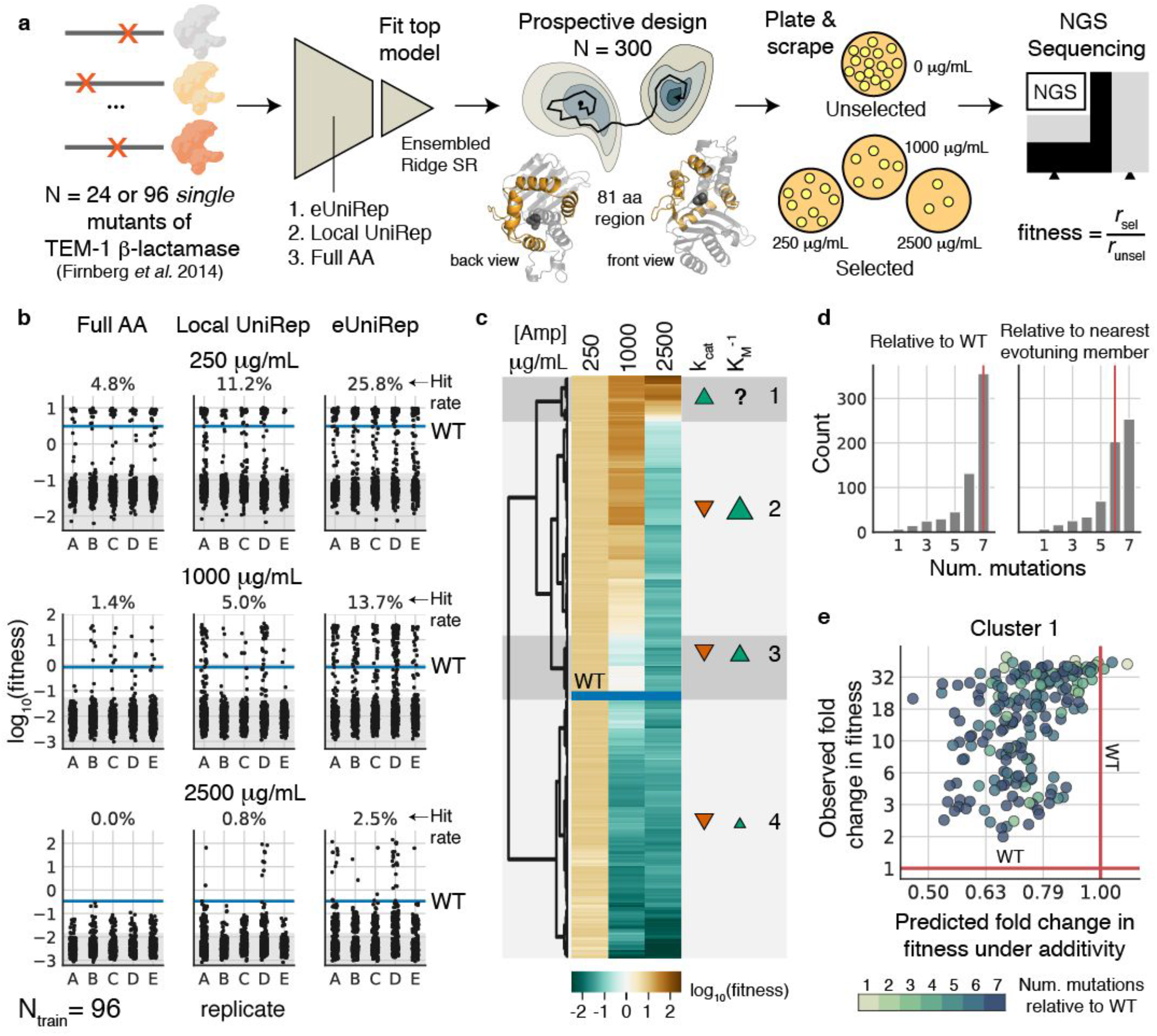
eUniRep enables low-N engineering of the enzyme TEM-1 β-lactamase using only single mutants as training data. **a)** Experimental workflow describing training mutant acquisition, sequence-to-function modeling, *in silico* directed evolution, and plate-based antibiotic selection combined with NGS sequencing to characterize designs. **b)** Low-N engineering results using N=96 training mutants for three different antibiotic selections. **c)** Heatmap illustrating log_10_(fitness) of all >WT eUniRep designs. Four clusters are annotated, and for each, likely changes to k_cat_ and K_M_^-1^ relative to wild-type are qualitatively shown. **d)** Bar plots illustrating the number of mutations of eUniRep designs to WT (left), and to the nearest member of the evotuning sequence set (right). **e)** Scatter plot of eUniRep Cluster 1 (highly >WT) designs illustrating observed fold change in fitness (relative to wild-type) vs predicted fold change in fitness under additivity.

Evotuning globally pre-trained UniRep was reproducible, and in 19 out of 20 replicates (95%), eUniRep enabled an overall 10 +/- 2% (95% CI) hit rate, defined as designs with activity greater than wild-type (>WT; eUniRep 1 & 2; Fig. 2b). For designs with 3 or fewer mutations, hit rates were 20%-65% and were substantially higher than those from error-prone PCR mutagenesis (Supplementary Fig. 4b,d), a typical starting point for directed evolution. Unexpectedly, maximal activity improvements *—* nearly 10x wild-type *—* were observed for designs containing 3-7 mutations, even though they had lower hit rates (5-25%). This reflects a risk-reward trade-off that eUniRep can exploit that would be challenging to achieve with directed evolution (Supplementary Fig. 4a,c).

**Figure 4.**
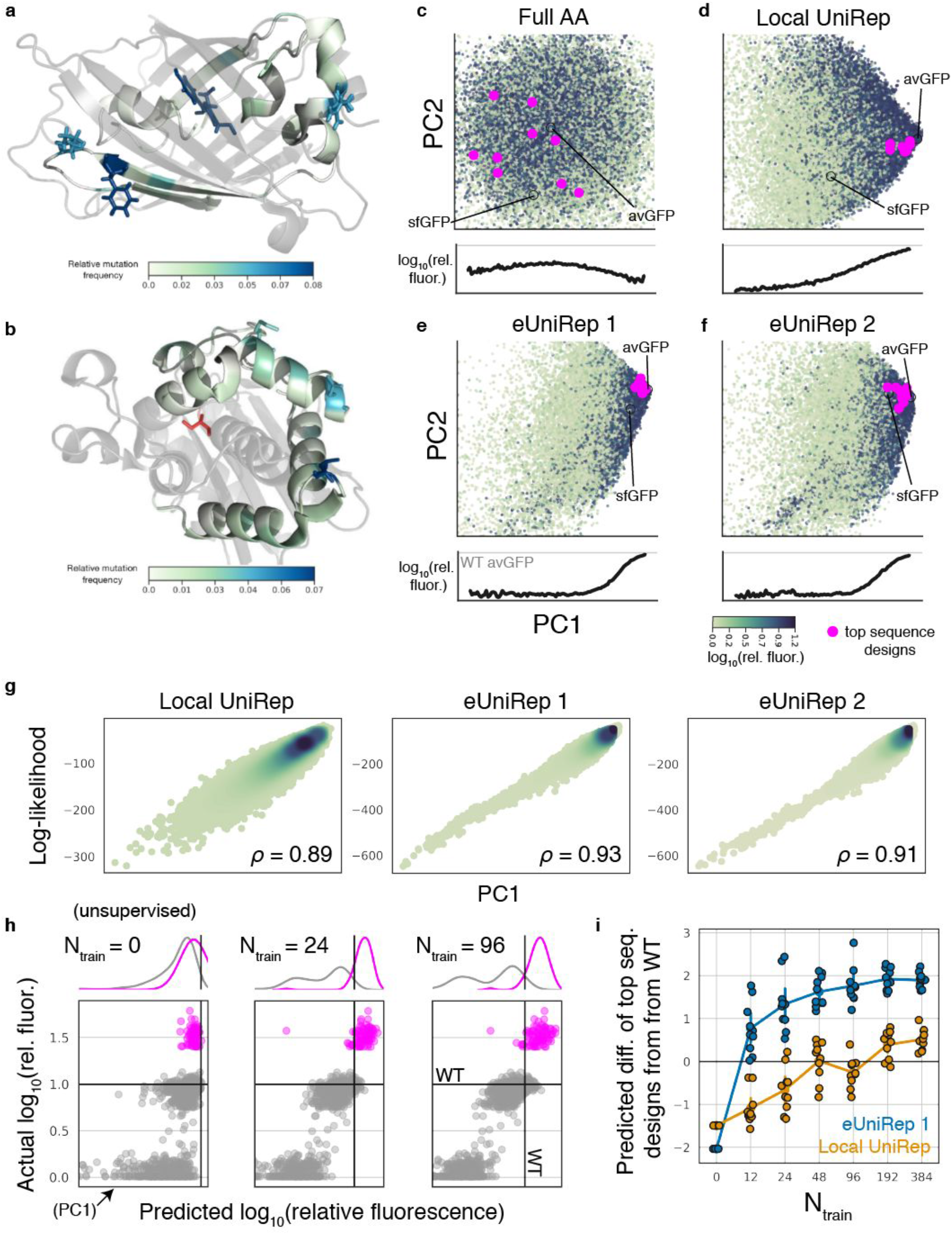
eUniRep designs are structurally non-trivial and require both unsupervised training and low-N supervised training to discover >WT variants. **a)** Structural visualization of avGFP (PDB: 2WUR). Mutations colored by relative frequency in >WT designs. Top 3 residues by mutation count shown as sticks. Chromophore colored by count of mutations made to any of the chromophore residues. **b)** As in **a)** but for the TEM-1 β-lactamase structure (PDB: 1ZG4), where the catalytic serine (S70) is highlighted in red. PCA of **c)** Full AA, **d)** Local UniRep, **e)** eUniRep 1 and **f)** eUniRep 2 representations of sequences from the local fitness landscape of avGFP, colored by log_10_(relative fluorescence). Magenta points show the top 10 sequence designs produced by each model. Below each plot, log_10_(relative fluorescence) as a function of PC1, Pearson *r* = 0.02 (Full AA), *r* = 0.52 (Local UniRep), *r* = 0.52 (eUniRep 1), *r* = 0.51 (eUniRep 2). **g)** Sequence log-likelihood vs PC1 for Local UniRep (left), eUniRep 1 (middle), and eUniRep 2 (right) with spearman correlations noted. **h)** Scatter plots of actual vs predicted log_10_(relative fluorescence) ordered by varying amounts of supervision. N_train_ = 0 corresponds to a purely unsupervised case, and so the x-axis corresponds to PC1. Grey circles are examples from the training distribution from which low-N training mutants are sampled. Magenta points represent the top 82 designed GFP sequences. Kernel density estimates of each population are shown above each scatter plot. **i)** Jitter plot depicting the degree to which top sequence designs can be differentiated from wild-type on the basis of predicted activity as a function of the number of low-N training mutants used (Methods). At a given N_train_, each datapoint represents a prediction replicate, which involves an independently sampled low-N training sequence set.

Repeating prospective design while constraining *in silico* evolution to a 7 mutation trust radius improved eUniRep’s overall hit rate to 18% without loss of quantitative fluorescence (Supplementary Fig. 5). Based on these numbers, “24-to-24 design” appeared tractable, where the characterization of just 24 training mutants and 24 optimized designs would be sufficient to observe a >WT design 1.8 +/- 0.8 (95% CI) times (Supplementary Fig. 6). By contrast, prospective design on Full AA or Local UniRep was inconsistent and only enabled ∼0% and ∼2% hit rates, respectively, highlighting the importance of both global and local unsupervised training.

We clonally validated our best designs and compared them to sequences produced by ancestral sequence reconstruction (ASR)^43,44^ and consensus sequence design^45,46^ (Methods). While both consistently provided >WT variants, eUniRep designs were substantially more functional (Fig. 2c). Several, in fact, were on par with superfolder GFP (sfGFP; Fig. 2c), which is the result of a multi-year engineering effort that started with avGFP and benefits from mutations outside of our design window. Importantly, eUniRep designs were diverse and occupied a unique region of sequence space, different from evotuning, ASR, and consensus sequences (median minimum number of mutations = 5, Fig. 2d).

Importantly, eUniRep’s design performance could not be explained by a simple tendency to guide search toward the evotuning or low-N training sequences. First, the vast majority (>99%) of evotuning sequences were less than 28% sequence similar to avGFP (>170 mutations; Supplementary Fig. 7a-b). Furthermore, of all mutations present in >WT eUniRep designs, approximately 25% were novel — defined to be neither found among the evotuning sequences nor the low-N training sequences (Supplementary Fig. 8a). For the remaining 75% of shared mutations, abundance among evotuning or low-N training sequences was a poor predictor of abundance among >WT eUniRep designs (Spearman *ρ* = -0.24). Finally, 89% of all >WT eUniRep designs contained at least one novel mutation, with many of the most active designs containing 33-66% novel mutations (Supplementary Fig. 8b). As these analyses only consider simple “first-order” mutational overlap, they provide a lower-bound on non-triviality. Indeed, due to epistasis, even recombining existing mutations among evolutionary homologs to produce functional proteins is a difficult challenge^47^.

### Low-N engineering of the enzyme TEM-1 β-lactamase

We next challenged our approach to generalize to the enzyme TEM-1 β-lactamase and optimize protein function training only on single mutants, which lack epistatic information^48^. Not only is this an arduous task due to the essential role of epistasis in proteins^49,50^, but also TEM-1 β-lactamase is dissimilar to avGFP both evolutionarily (Eukaryotic vs. Prokaryotic) and functionally (fluorescence vs. hydrolysis). Additionally, unlike GFP, our measure of TEM-1 β-lactamase function is only observable through organism level fitness (Methods), which is an indirect, endpoint measure that depends on the activity of other proteins (e.g. peptidoglycan forming DD-carboxypeptidases and peptidoglycan transpeptidases). Finally, we note that low-N engineering is particularly desirable for enzyme biocatalysts^18^, of which β-lactamase is a model. Here, high-throughput assays are frequently intractable due to the difficulty of intracellularly reporting on enzyme activity.

We performed low-N optimization of TEM-1 β-lactamase fitness in 3 concentrations of the antibiotic ampicillin (250, 1000, or 2500 μg/mL) using single mutants as training data (Fig 3a; Methods; Supplementary Fig. 9)^48^. We designed a 81 amino acid region spanning four helices that straddle, but do not include the central helix bearing the catalytic serine, S70 (Fig. 3a). Designs were proposed with a 7 mutation trust radius (Methods). As done with GFP, we generated 300 designs for each N_train_ and representation model and replicated this process 5 times.

eUniRep consistently enabled a 5-10x and 2-3x higher hit rate than Full AA and Local UniRep, respectively (Fig. 3b). eUniRep’s relative performance improved to a 5-9x gain over Local UniRep for training sets of size N=24 (Supplementary Fig. 10), and except at the most stringent antibiotic concentration, eUniRep’s performance was robust and consistent across training sets.

Importantly, eUniRep designs were diverse both in function and in sequence (Fig. 3c-d). A hierarchical clustering of log-fitness profiles and a qualitative approximation of Michaelis-Menten kinetics revealed >WT eUniRep designs could be grouped into four clusters, consistent with changes in k_cat_ and K_M_ (Fig. 3c; Methods). As observed for GFP, eUniRep >WT designs were significantly diverged from wild-type (median number of mutations = 7) and from any evotuning set sequences (median minimum number of mutations = 6) (Fig. 3d).

We note that the majority (>89%) of evotuning sequences had less than 28% sequence identity to wild-type TEM-1 β-lactamase (>204 mutations; Supplementary Fig. 7c). On average, 18% of the mutations found in eUniRep >WT designs were novel (Supplementary Fig. 11a), and of those that were not, abundance among the evotuning sequences was not a strong predictor of abundance among designs (Spearman *ρ* = 0.1). Additionally, 97% of >WT designs contained at least 1 novel mutation, and many of the most active designs contained 30-70% novel mutations (Supplementary Fig. 11b). Therefore, as with GFP, it is unlikely that the higher hit rates of eUniRep are explained by a simple tendency to guide search toward sequences in the evotuning or low-N training sequence sets.

Notably, despite being generated from single mutant training data, eUniRep’s >WT designs were epistatically non-trivial (Fig. 3e). For Cluster 1 designs, which were >WT in all antibiotic conditions, we calculated predicted fitness assuming each mutation contributed additively, and compared this to the experimentally observed fitness of the fully mutated design. Surprisingly, most of these designs were substantially >WT despite their prediction under additivity being loss-of-function (Fig 3e). Additionally, their *in silico* evolutionary trajectories were consistent with the navigation of a rugged, epistatic fitness landscape^51^ (Supplementary Fig. 12). These results suggest that via transfer of epistatic information from unsupervised learning, eUniRep can exploit epistasis even when no higher-order mutation combinations have been observed in the training data.

### Unsupervised training serves to guide search away loss-of-function sequences, while low-N supervision enables the discovery of >WT sequences

We next attempted to explain eUniRep’s unique ability to enable low-N engineering (Fig. 4, Supplementary Fig. 13-15). While mutations in eUniRep proposals and >WT designs were biased toward solvent-exposed residues, a substantial fraction (40% GFP and 28% β-lactamase) were targeted to buried positions including the avGFP chromophore (Fig. 4a, Supplementary Fig. 13). This suggested that eUniRep could make non-trivial, beneficial rearrangements to the hydrophobic core, which previous work has suggested is difficult^29^. Additionally, we observed that the most functional β-lactamase designs were not preferentially mutated near the catalytic serine (S70), which ran counter to the typical engineering heuristic of targeting mutations around the enzyme’s active site^19^. This result also suggested eUniRep can exploit non-local epistatic interactions (Fig. 4b, Supplementary Fig. 13). Unsurprisingly, eUniRep’s mutational preference could not be explained by first-order position-wise mutational tolerance, suggesting that eUniRep enabled more than consensus sequence design despite both methods drawing on evolutionary information (Fig. 2c, Supplementary Fig. 14).

Not finding a clear explanation for eUniRep’s performance among these structural and evolutionary analyses, we examined the eUniRep sequence representation. Strikingly, we found a strong correlation between its primary axis of variation (principal component 1; PC1) and protein function (Fig. 4c-f, avGFP Pearson *r* = 0.51, 0.52; Supplementary Fig. 16a, β-lactamase Pearson *r* = 0.44), which was not observed for PC1 of the Full AA representation (avGFP Pearson *r* = 0.02, β-lactamase Pearson *r* = 0.05). However, while PC1 could differentiate non-functional sequences from functional ones, it could not differentiate functional sequences with wild-type or greater levels of activity (Fig. 4c-f, Supplementary Fig. 16a). For example, our most active GFP designs and sfGFP had similar PC1 scores to wild-type avGFP (Fig. 4e-f). Further examination revealed that PC1 was highly correlated with sequence likelihood under each UniRep model, with the highest such correlations observed for eUniRep (Spearman *ρ* = 0.93 and 0.91 for eUniRep 1 and 2, respectively; Fig. 4g, Supplementary Fig. 16b). Given global unsupervised pre-training and evotuning of these models are performed on natural sequences, this suggests that the primary utility of unsupervised learning as performed here is to guide search away from unpromising sequences in the fitness landscape based on a (semantically meaningful) sense of their unnaturalness.

However, it also suggests that unsupervised training alone does not enable the discovery of better-than-natural variants. Indeed, we observed that only with low-N supervised learning could >WT designs be differentiated from those with wild-type or lower levels of activity (Fig. 4h-i, Supplementary Fig. 16c-d). Thus, we propose a two part model to explain eUniRep’s ability to enable low-N protein engineering: First, unsupervised learning greatly simplifies search by eliminating the vast majority of the non-functional fitness landscape on the basis of unnaturalness. “On top” of this information, supervised learning with a small number of low-N mutants then distills the critical information needed to discover better-than-natural variants.

## Discussion

This work is the first to demonstrate a generalizable and scalable paradigm for low-N protein engineering. By distilling information from both the global and local sequence landscape, we reproducibly leveraged N=24 random training mutants and one round of *in silico* screening into over 1000 novel >WT designs. This is the strongest case of generalization- and data-efficiency in machine learning guided protein function optimization to date (Supplementary Fig. 17). Additionally, our two part mechanism to explain this performance provides context for and extends previous unsupervised protein function modeling and design work. While unsupervised methods trained on natural sequence data perform well at predicting or avoiding loss-of-function variants during modeling and design, they have also been unable to reliably model or design better-than-natural variants^38,39,41,47,52^. Our findings suggest that a small amount of labeled data and additional supervised learning on top of unsupervised pre-training may be necessary to find enhanced variants.

We took advantage of robust, high-fidelity multiplexed assays to extensively characterize our approach on avGFP and TEM-1 β-lactamase. While low-N design is intended for proteins where such assays are not available, both proteins have a rich history of being studied or engineered with them. As such, we consider existing >WT variants to be a high bar. Here, with just 24 random mutants of avGFP as training data, we designed novel FPs that rivaled sfGFP, the product of many years of high-throughput, high-fidelity protein engineering.

Nevertheless, unlike GFP and TEM-1 β-lactamase, most proteins do not have assays that are both high-throughput and high-fidelity. In many therapeutic and industrial projects, high-fidelity experimental measurements of endpoint functions, like crop yield or biologic efficacy, are scarce and come at the end of long test cycles. In theory, generating high-throughput proxy assays of these endpoints should improve engineering success rates. However, empirically this is often not the case as evidenced, for example, by Eroom’s law in drug development^13,15^. Here efforts to use high-throughput proxy assays for the endpoint in question may in fact generate worse candidates for later-stage development^13,15^ by over-optimizing a biased metric^53^. Taken together, this suggests generalizing from low-N high-fidelity measurements may be more important than learning from high-N low-fidelity measurements.

Indeed, several previous efforts successfully engineered valuable proteins using high-fidelity assays and low-N design^19,23,24,54–58^. However, these (semi-)rational protein engineering approaches intensively rely on hand-crafted structural or (co-)evolutionary priors to narrow the search space of potential mutations^8,19,59,60^. Additionally, they often require expert judgment to learn from data, which may include modifying energy functions for biophysical design^61^, and iteratively designing and testing structure-guided mutation combinations^19,62–65^. Together these modeling and design choices introduce biases that could manifest as a mismatch between optimization metric and endpoint. By contrast, UniRep and our low-N approach are paradigmatically empirical and sequence-based, improving with the exponential growth of sequence databases to minimize bias^25^, and leaving open the possibility of discovering new principles of protein folding and activity that extend beyond our current mental models. Indeed, when combining data-driven digital fitness landscapes with *in silico* evolution to both measure well and search far, we find there may be surprising diversity and function in the vastness of sequence space.

## Acknowledgements

We thank Mohammed AlQuraishi, Chris Bakerlee, Anush Chiappino-Pepe, Aleksandra Eremina, Kyle Fish, Sager Gosai, Xiaoge Guo, Eric Kelsic, Sri Kosuri, Pierce Ogden, Sam Sinai, Max Schubert, Amaro Taylor-Weiner, David Thompson, and Aaron Tucker for feedback on earlier drafts of this manuscript. We thank members of the Esvelt and Church labs for valuable discussion. S.B. was supported by an NSF GRFP Fellowship under grant number DGE1745303. G.K. was supported by a grant from the Center for Effective Altruism. E.C.A. was supported by a scholarship from the Open Philanthropy Project. This material is based upon work supported by the U.S. Department of Energy, Office of Science under Award Number DE-FG02-02ER63445. Computational resources were, in part, generously provided by the AWS Cloud Credits for Research Program and Lambda Labs, Inc.

## Author contributions

S.B., G.K., E.C.A. conceived the study. S.B. performed wet-lab experiments and managed data. S.B., G.K., E.C.A. performed machine learning modeling and data analyses. K.E. and G.M.C. supervised the project. S.B., G.K., E.C.A. wrote the manuscript with help from all authors.

## Data and Code availability

Code for UniRep model training and inference with trained weights along with links to all necessary data is available at https://github.com/churchlab/UniRep.

## Methods

### Evolutionary fine-tuning (evotuning)

We reasoned that by fine-tuning UniRep’s existing knowledge of all protein sequences to the evolutionary neighborhood of the target sequence (evotuning), we may be able to reduce the prohibitive data requirements of supervised deep learning and thereby enable low-N design. Indeed, impressive gains in data-efficiency have been obtained through similar means in other machine learning domains including vision^27,66,67^ and language^66,68,69^. We began with model weights that had been globally pre-trained on UniRef50 as described previously^25^. To evotune, we select a subset of public sequences which are closer to the target protein, and then finetune the globally pre-trained weights on the UniRep mLSTM model on this local sequence neighborhood.

For avGFP, we used the same evotuned weights as previously described, called eUniRep 1^25^ above, and additionally repeated the evotuning process to ensure its robustness. As with eUniRep 1^25^, the avGFP target sequence together with a selection of related fluorescent proteins was jackHMMer searched^70^ until convergence. Edit distance was computed between the search result sequences and the avGFP target sequence. The sequence set was filtered for length (kept all <500 amino acids) and Levenshtein distance from avGFP (kept all <400), and sequences with non-standard amino acids were removed, yielding 79,482 sequences. We note that this number is larger than the 32,225 sequences used to train eUniRep 1 obtained in reference^25^. The difference is due to the stochasticity of JackHMMER and updates to the JackHMMER web server between runs for eUniRep 1 and eUniRep 2, as well as running JackHMMER to convergence for eUniRep2. We note that the downstream design performance enabled by these two evotuning models was similar despite this 2x difference in the number of sequences in the dataset.

To determine when to stop training, we selected a 10% “out of distribution set” by sampling each sequence with a probability proportional to the 4th power of the edit distance. A 10% in-distribution set was selected uniformly randomly. We initialized the weights of the 1900 dimensional UniRep mLSTM with the globally pre-trained weights and trained for 13,500 iterations with early stopping^71,72^, until the outer validation set loss began to increase. This model was used to produce the representations for eUniRep 2 as named above.

The evotuning for TEM-1 β-lactamase proceeded similarly, seeding the jackHMMer search with the wild-type TEM-1 β-lactamase together with related beta-lactamase sequences. The results were filtered for length (<600 amino acids) and Levenshtein distance from TEM-1 β-lactamase (<286) and sequences with non-standard amino acids were removed yielding 76,735 results. Training, initialized with the global weights as above, proceeded for 13,500 iterations.

For Local UniRep, we used the same dataset and training procedure as above, but instead of using the globally pre-trained UniRep weights as initialization, we generated a random weight initialization from the same distribution that was used to initialize the original UniRep model. This is analogous to retraining the original UniRep model but just on the local sequence landscape, leading to the name Local UniRep.

### Retrospective experiments for low-N engineering

The purpose of our retrospective experiments was to evaluate the possibility of low-N engineering. Toward this end, we tested the abilities of different sequence-to-function models meaningfully generalize in terms of predictive performance from a “local” region of the fitness landscape to more “distant” regions using only a small number, N, of (sequence, function) pairs from the local fitness landscape.

Our retrospective experiments took the following steps:

1. Dataset creation and processing. Here we established three datasets whose generation and/or processing is described in detail below and whose properties are summarized in Supplementary Figure 1:
  a. “Sarkisyan”, which is comprised of functionally characterized sequences from the local fitness landscape of avGFP. This dataset was publicly available and was processed from Sarkisyan *et al*. (2016)^42^. In our experiments, this dataset was used for sampling training sequences.
  b. “SynNeigh”, which is comprised of functionally characterized sequences from the local fitness landscape of sfGFP and the local fitness landscapes of related variants of sfGFP that were obtained through simple ML guided exploration strategies. Thus, this dataset represents a collection of many local fitness landscapes for different avGFP’s. This data was generated from variants obtained from Biswas *et al*. (2018)^29^, and will be made publicly available upon peer-reviewed publication. In our experiments, this dataset was used for evaluating generalization.
  c. “FP Homologs”, which is comprised of functionally characterized sequences from the global fitness landscape of known *Aequorean* fluorescent proteins. This dataset was generated by molecularly shuffling the DNA of 65 extant *Aequorean* FPs, and thus represents a global, albeit sparse sampling of the global fitness landscape significantly beyond that explored in the local fitness landscape of avGFP (Sarkisyan). This dataset was generated and processed for this work, and will be made publicly available upon peer-reviewed publication. In our experiments, this dataset was used for evaluating generalization.
2. Each dataset was then randomly split three ways to produce Splits 0, 1, and 2. Model prototyping and evaluation as described in subsequent steps below was entirely performed on Split 0. After prototyping, a final list of models, hyperparameters, and procedural parameters were fixed and performance of each approach was evaluated on Split 1, the results of which are reported in Supplementary Figure 2. Split 2 was used for all prospective experiments as reported in the main text.
3. On Split 0, we systematically evaluated the impact of several factors on generalization. We defined good generalization to be accurate rank ordering of sequences in a generalization set, such that if we were to select the top ranked sequences for experimental characterization, they would be highly functional. The factors examined are as follows:
  a. Number of training sequences (N).
  b. Acquisition policy - This defines how the N training sequences are selected. A complete list of policies and their descriptions are below.
  c. Sequence representation - This defines how the amino acid sequence is numerically encoded to the top-model. Full AA or eUniRep are examples of encodings. A complete list of representations and their descriptions are below.
  d. Top model - This is a simple, low-parameter supervised model that is trained on training sequence representations to predict quantitative function. Ridge regression is an example top model. A complete list of top models examined and their descriptions are below.
4. Once we were able to determine how these variables affected retrospective generalization, especially in low-N settings, we fixed a final list of N training sequences, sequence representations, top-models, and reporting criteria and reproduced the retrospective experiments again on Split 1. This was to ensure we did not overfit to Split 0. A summary of these results are reported in Supplementary Figure 2.

Retrospective experiments result summary -

Supplementary Figure 2 summarizes the results of our retrospective generalization experiments, where the task is to rank order members of the generalization set such that if we were to select the top 96 for characterization as many as possible should be >WT “hits”. To contextualize performance, this metric can be normalized as a ratio to the performance obtained by a random ordering of generalization set members.

Sequence representation was the most influential variable that affected performance. One-hot Full AA, Doc2Vec^73^, UniRep (globally trained, but not evotuned) generally did not show improvements over random for any size training set from the local fitness landscape of avGFP (Sarkisyan). By contrast, evotuned models showed a greater than 20x performance gain over random when generalizing to members of SynNeigh and x performance gain over random when generalizing to members of FP Homologs. In particular, eUniRep 1 and eUniRep 2 were superior to Local UniRep, which lacks knowledge of global sequence space, showing highly data-efficient performance with as few as N=8 training sequences (Supplementary Fig. 2).

Choice of top-model played a less significant but nonetheless important role. In particular, we noticed a marked performance difference between L1- (Lasso/LARS) and L2-penalized (Ridge) top models, with L2 variants performing substantially better. We suspect that this is likely because the meaningful information contained in the mLSTM representations are entangled and hence the representation as a whole is non-sparse. This violates the assumptions of L1 penalized regression. Among L2 models, we noticed that choosing a more stringent regularization with the same (statistically) inner cross-validation performance gave a slight performance gain (Ridge SR). Finally, ensembling this approach (Ens Ridge SR) neither hurt nor improved performance, but gave us an empirical uncertainty estimate.

Interestingly, how training sequences were acquired did not matter much (data not shown). For real-world technical simplicity we therefore chose to acquire training points randomly from the output of error-prone PCR or single-mutation deep mutational scans. Additionally, the models that worked the best (eUniRep powered models) were surprisingly robust to the number of training points sampled (N= 8, 24, or 96), which are all small enough that they can be feasibly collected for a variety of proteins and applications.

### Dataset creation

Three datasets were used for our retrospective low-N engineering experiments. The Sarkisyan dataset was also used for the prospective design experiment illustrated by Figure 2 in the main text. A detailed description of their generation and/or processing follows:

#### Sarkisyan -

This dataset was obtained from Sarkisyan *et al*. (2016)^42^; it is publicly available. Briefly, the authors used error-prone PCR to mutate wild-type avGFP, and then measured the fluorescence of approximately 50,000 variants using FlowSeq in a manner similar to how it was performed in this work (see “FlowSeq” section). We further processed their dataset by:

1. Min-max scaling log_10_(relative fluorescence) values according to the formula, (x - min_val)/(wt_val - min_val), where min_val is the fluorescence of the least fluorescent sequence and wt_val is the fluorescence of the wild-type sequence. Thus, after transformation wild-type fluorescence corresponds to a value of 1, whereas an entirely non-functional sequence has fluorescence 0. This min-max scaling was performed to ensure consistency with the other datasets.
2. Random splitting of the dataset into 3 splits as described above.

The distribution of transformed fluorescence values, edit distances (number of mutations) to avGFP, and edit distances between members of this dataset are shown in Supplementary Figure 1a.

#### SynNeigh -

The purpose of this dataset was to serve as a generalization set to evaluate model generalizability. This dataset was generated from variants discovered in Biswas *et al*. (2018)^29^. Here the authors used a variety of simple machine learning guided approaches to propose diverse but functional sequence variants of sfGFP. This included model guided exploration under a three layer fully connected feed-forward neural network and under a composite-residues neural network. The goals of these explorations were varied, and included attempts to improve fluorescence, diversify the sequence while maintaining function, and to diversify the sequence while maintaining function while only mutating combinations of otherwise difficult to singly mutate residues. In total, 286 “parent” variants were proposed in this manner.

In this work, after pooling plasmid DNA for all 286 parent variants, we performed error-prone PCR (GeneMorph II Random Mutagenesis Kit, Agilent Technologies) over the full length of the GFP gene aiming for an average of 2 mutations per template. This library was cloned and transformed into DH5α *E. coli* (see “Library Cloning and transformation” section), with an estimated library size of 150,000. The relative fluorescence of each variant in the library was then measured with FlowSeq (see “FlowSeq” section). In total, we obtained high-quality fluorescence measurements for 104,285 variants.

Because many of the 286 parent variants are highly functional and we were mostly measuring minorly mutated variants thereof, much of the dataset is comprised of functional variants. In practice, in low-N engineering our task is to find rare high-functioning sequences among a sea of non-functional sequences in the distant or non-local fitness landscape. To better incorporate this intuition into our retrospective experiments we therefore filtered out variants with intermediate fluorescence (>= 0.7 and <= 1.5), leaving only non-functional and highly functional variants. After filtering, we retained 52,512 variants, 52,416 of which were non-functional and 96 of which were highly-functional.

This final dataset, which we refer to as “SynNeigh”, was min-max scaled (using avGFP fluorescence as wt_val) as above and then split into 3 parts. Because all measured variants were derivatives from one of the 286 parents, the three-way split was created by first randomly splitting the 286 parent variants three ways and then assigning derivative variants to one of the three splits according to the parent variant to which they had the fewest mutations.

#### FP Homologs -

The purpose of this dataset was to serve as an additional generalization set to evaluate model generalizability, and was generated for this work. While SynNeigh is inherently “centered” around sfGFP, and samples several local fitness landscapes densely, FP Homologs sparsely samples the global fitness landscape of known *Aequorean* FPs.

To accomplish this, we first mined an October 2018 download of the FPBase database^74^ for FP sequences of *Aequorean* origin. Of these 132 sequences, 70 were mutually different by at least three mutations. After manual curation of these 70 sequences, which involved stripping away His-tags, and manually adjusting the N- and C-termini of the sequences which were sometimes modified for crystallization purposes, 65 sequences remained that were mutually different by at least one mutations. The median and maximum number of amino acid mutations between these 65 “parent” sequences was 15 and 63, respectively. Note, the full length of each sequence was 238 amino acids. These parents also encompassed a variety of spectral properties, with some of them fluorescing blue or yellow in addition to green. Nucleotide sequences of these 65 parents were obtained by choosing an *E. coli* codon optimization and were subsequently ordered as separate Gene Fragments from Twist Biosciences.

Each Gene Fragment was cloned into DH5α *E. coli* individually using Golden Gate assembly, and the coding sequence and spectral phenotypes were individually confirmed. Plasmid DNA for each parent was mini-prepped (Qiagen) and all parent plasmid DNA was subsequently pooled. To generate a sparse, but broad sampling of sequences in the global fitness landscape spanned by these parents, we performed DNA shuffling^75^ followed by error-prone PCR (GeneMorph II Random Mutagenesis Kit, Agilent Technologies). The DNA shuffled and error-prone PCRed library is hereafter referred to as the “Shuffled Library.”

Because Shuffled Library contained mutations throughout the full length of the FP gene, it was not immediately compatible with our FlowSeq protocol, which cannot sequence more than a 600 bp amplicon. We next therefore performed a “stitching PCR”, where we added a random 20 bp DNA barcode (BHVDBHVDBHVDBHVDBHVD) to 3’ end of the DNA in both the Parent Pool and Shuffled Library, after the stop codon of the gene. The now barcoded Parent Pool and Shuffled Library were separately cloned and transformed into DH5α *E. coli* (see “Library Cloning and Transformation” section), with an estimated library size of approximately 100,000 members. Through simulation we confirmed it would be overwhelmingly statistically likely that one barcode would “point to” just one template and not more. Barcodes did not affect translation, but were likely transcribed. We nonetheless assumed this would have a negligible impact on the expression level of the resulting protein.

We next spiked in the transformed Parent Pool at 0.5% into the transformed Shuffled Library and performed FlowSeq (see “FlowSeq” section). Because this pool contains a collection of spectrally diverse variants, we excited with two different laser combinations (488 nm only, 405 nm + 488 nm) and sorted in four different emission channels (FL1=450/50 3 bins, FL2=525/50 8 bins, FL3=600/60 6 bins, and FL4=665/30 2 bins). Instead of sequencing the coding region, we sequenced the 20 bp barcode. Barcode sequencing was done using a 2 × 75 bp NextSeq mid-output sequencing run.

Examining a heatmap of variant log-abundances across all samples, we observed clear structure indicating groups of variants that were clearly enriched or depleted from sort bins representing different fluorescence intensities under different excitation (lasers) and emission (filters) conditions. However, we also observed what we suspected to be higher frequency noise in which certain variants would be abundant in one condition but would have zero counts in a highly related condition. We suspected this was an artefact of under-sorting and possibly under-sequencing our library. To remedy this, we performed imputation of these missing measurements with MAGIC^76^, which was originally developed to perform the same kind of imputation for drop-out measurements in single-cell RNA-seq data. We confirmed imputations were likely high-fidelity by artificially dropping out measurements of high-confidence variants (the highly abundant parent sequences) and examining the accuracy of their imputed values (Pearson *r* = 0.89). Considering these imputed counts as “final”, we proceeded with fluorescence inference as we would for a normal FlowSeq experiment. At this point we obtained log_10_(relative fluorescence) values associated with each barcode, and for consistency, specifically used those associated with 405 nm + 488 nm excitation and emission in FL2 (525/50).

In order to determine the identity of the variant each barcode represented, we performed long-read amplicon sequencing. The sequenced amplicon included both the coding sequence of the FP as well as the 3’ barcode. Two independent PacBio Sequel II runs were performed. The first was of the Parent Pool and Shuffled Library (input into FlowSeq). The second was of all functional members of the Parent Pool and Shuffled Library, which was deemed to be all variants that didn’t sort into the non-functional bin during the FACS step of FlowSeq. The second was done to increase the chances we could successfully decode barcodes for functional library members.

After performing a number of sanity checks, we could reliably associate barcodes with their respective FP variants. The number of instances a given barcode pointed to multiple variants that were not explainable by sequencing noise was extremely low (<1e-2%). In total, we could make 40,581 high-confidence barcode associations, representing 37,582 unique variant sequences. In total, these 37,582 variants (and their 40,581 associated barcodes) accounted for 58% percent of the NextSeq barcode sequencing data after basic processing (read pair merging, amplicon extraction, and basic length filtering on the barcodes). This suggested, that while it’s likely a small to moderate size of transformed library might have been missed using this barcode association procedure, we could still capture a large fraction of it.

To make the generalization task more challenging we further filtered this data to include only parents that were highly functional (10x brighter than avGFP) and variants that beared any of their sequence. To do this, we first identified a set of 16 parent sequences that were highly functional (>10x brighter than avGFP) and confirmed their qualitative improvement over avGFP from the literature. We then analyzed the protein sequence of every variant and assigned any variant with any subsequence that could be unambiguously attributed to one of these 16 parents to be in the filtered list of variants. 27,050 variants met these criteria.

Finally, as done for SynNeigh, we removed variants with intermediate fluorescence, min-max scaled the fluorescence values as above, and split the data randomly into three splits.

### Acquisition policies

We considered several acquisition policies for sampling training set (sequence, function) pairs. These could be broadly classified into three categories, sequence-only, structural, and evolutionary based on the primary source of information they need. For sequence-only methods, we considered randomly sampling mutants from the output of error-prone PCR and randomly sampling single mutants (e.g. as the output of a deep mutational scan). For structural and evolutionary approaches we considered several policies that would sample mutations based on their structural and evolutionary conservation properties in order to build epistatically dynamic training sets. We found the sequence-ony policies of random sampling from error-prone PCR or from single mutants to be as performant as structural and evolutionary policies.

### Sequence representations

We considered several different ways to convert sequences into a numerical representation suitable for use in supervised modeling.

1. Full AA - one-hot encoding of the full amino acid sequence is a simple representation method that exactly represents the information contained an amino acid sequence; no more, no less. Procedurally, to one-hot encode a sequence of length L, a 20 x L matrix, O, is constructed such that O[i,j] = 1 if amino acid i occurs in position j of the sequence (for some predetermined ordering of the 20 amino acids). The final encoding of the sequence is a “flattened” or “unrolled” version of O, that is a vector of dimension 1 x (20*L).
2. Doc2Vec - Here we use a previously state-of-the-art approach for representing protein sequences^73^, based on the popular Doc2Vec natural language processing paradigm for generating vector representations of entire documents^77^. In previous work where we developed UniRep, we compared extensively to this Doc2Vec-for-proteins approach^25^.
3. UniRep - The sequence representation obtained from the globally trained (on UniRef50) UniRep mLSTM. Specifically, the representation is the average hidden state taken across the length of the sequence as reported in Alley *et al*. (2019)^25^. We also refer to this representation as “avg_hidden.”
4. Local UniRep - The avg_hidden representation obtained from training a randomly initialized mLSTM whose architecture is the same as UniRep on the same local sequence dataset used for evotuning.
5. eUniRep - The avg_hidden representation obtained from Evotuning the UniRep mLSTM that has already been globally trained on UniRef50. The additional suffixes of “1” or “2” refer to replicates of the Evotuning process.

### Top models

We considered several top models. Though in principle any supervised model could be used here, for the purposes of low-N engineering, we reasoned that only simple low-parameter models would be reliably fit and have a lower risk of overfitting. Additionally, if the sequence representation is truly semantically rich, then only a simple top model should be needed to make accurate quantitative predictions about function. We therefore restricted our attention to single-layer models, i.e. various forms of linear regression:

1. Lasso-Lars - This is L1-penalized linear regression implemented using the Least Angle Regression algorithm^78^. We used the Python sklearn.linear_model.LassoLarsCV implementation to perform 10-fold cross-validation (on the input training data) to select a level of regularization (the parameter “alpha”) that minimizes held-out mean squared error. The schedule of regularization strengths is known up-front by the LARS algorithm.
2. Ridge - This is L2-penalized linear regression. We used the Python sklearn.linear_model.RidgeCV implementation to perform 10-fold cross-validation (on the input training data) to select a level of regularization (the parameter “alpha”) that minimizes held-out mean squared error. The schedule of regularization strengths was set to be logarithmically spaced from 1e-6 to 1e+6. Features were normalized up-front by subtracting the mean and dividing by the L2 norm.
3. Ridge SR - This is the same as the “Ridge” procedure above, except that we additionally perform a post-hoc “sparse refit” (SR) procedure. The “Ridge” top model above chooses a level of regularization that optimizes for model generalizability if the ultimate test distribution (i.e. distant regions of the fitness landscape) resembles the training distribution. However, this is not likely the case. Therefore, we perform a post-hoc procedure to choose the strongest regularization such that the cross-validation performance is still statistically equal (by t-test) to the level of regularization we would select through normal cross-validation. This procedure selects a stronger regularization than what would be obtained using the “Ridge” procedure as defined above.
4. Ensembled Ridge SR - This is the same as the “Ridge SR” procedure above, except that the final top model is an ensemble of Ridge SR top models. The ensemble is composed of 100 members. Each member (a Ridge SR top model) is fit to a bootstrap of the training data (N training points are resampled N times with replacement) and a random subset of 50% of the features. The final prediction is an average of all members in the ensemble. The rationale for this approach is that it is based on consensus of many different Ridge SR models that have different “hypotheses” for how sequence might influence function. Differences in these “hypotheses” are driven by the fact that every bootstrap represents a different plausible instantiation of the training data and that every random subsample of features represents different variables that could influence function.

### Training datasets for prospective low-N engineering

For prospective design of GFP, we relied on sampling random N=24 or N=96 sized subsets from the Sarkisyan dataset (see dataset descriptions in “Retrospective experiments for low-N engineering” above). This corresponded to virtually picking random mutants (e.g. colonies) from error-prone PCR generated library. This would be straightforward to implement experimentally, and indeed, error-prone PCR is a common starting point for many protein engineering efforts. A shortcoming of error-prone PCR is that because only a few nucleotide changes (usually at a rate of 0.1-0.5%) are made per gene, it is difficult to observe amino acid substitutions that require multiple mutations to the same codon. However, it is a simple and tunable way to sample higher-order mutation combinations.

For prospective design of TEM-1 β-lactamase, we relied on sampling random N=24 or N=96 sized subsets from the single-mutation scanning mutagenesis (deep mutational scan) dataset generated in Firnberg *et al*. (2014)^48^. Briefly, they performed scanning mutagenesis of the *E. coli* TEM-1 β-lactamase protein and profiled the activity of 95.6% (5,212/5,453) of single amino acid substitutions. Unlike the output error-prone PCR, scanning mutagenesis as performed here can explore any amino acid substitution. However, higher order mutation combinations were not explored. The authors used a tunable bandpass genetic selection assay^79^ measure the resistance of a variant to different concentrations of ampicillin, up to 1,024 μg/mL. The output of their assay was highly correlated with the minimum inhibitory concentration of ampicillin at which a variant can no longer confer resistance. We note that this is a different measure of fitness than we use in this work, which is based on log-fold enrichments. Nevertheless, we would expect a gain/loss-of-function variant in their system to be gain/loss-of-function in ours and so we felt it was a suitable pool of training mutants for our prospective design experiments.

### Prospective design: sequence proposal via *in silico* directed evolution

We wished to use an algorithm that would on average seek more functional variants, but was not deterministically forced to do so. We therefore utilized a Metropolis-Hastings Markov-Chain Monte Carlo algorithm to stochastically sample from the non-physical Boltzmann distribution defined by:

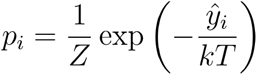

Where *ŷ*_*i*_ is the model predicted fitness for sequence *i,k*,is a constant that was set to 1,*T* is the temperature, and *Z* is an unknown normalization constant.

Our *in silico* directed evolution algorithm was as follows:

1. Input:
  a. An initial sequence
  b. A sequence-to-function model that predicts an amino acid sequence’s quantitative function, or fitness.
  c. Temperature, *T*.
  d. Trust radius: the number of mutations relative to wild-type allowed in proposed designs.
2. Initialize: set state sequence,*s*, equal to a provided initial sequence.
3. Propose a new sequence,*s*\*, by randomly adding *m* ∼ Poisson (μ − 1) + 1 mutations to *s*.
4. Accept proposal and update the state sequence,*s* ← *s*\*, with probability equal to 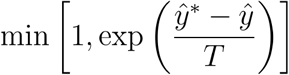, where *ŷ*\* and *ŷ* are the predicted fitness of the proposed sequence and state sequence, respectively. Otherwise, reject the proposal (and keep the state sequence as is). Note that if the sequence proposal has more mutations than the input trust radius, its predicted fitness is set, post-hoc, to negative infinity thereby forcing rejection of the proposal.
5. Iterate steps 3 and 4 for a predetermined number of iterations.

For the prospectively designed GFP and TEM-1 β-lactamase libraries, for a given sequence-to-function model (the combination of sequence representation method and a low-N trained top-model), 3500 evolutionary trajectories were run in parallel for 3000 iterations. The initial sequence for each trajectory was obtained by making Poisson(2)+1 random mutations to the wild-type sequence. The sequence proposal mutation rate,, for each trajectory was set to be a random draw from a Uniform(1, 2.5) distribution.

We investigated a number of different temperature parameters spanning six orders of magnitude. We found that for GFP and TEM-1 β-lactamase models a temperature of 0.01 gave good trajectory behavior. We qualitatively ascertained this by visualizing how predicted fitness varied across the trajectory. High temperatures, which increases acceptance probabilities, produced overly explorative trajectories that mostly dwelled in low predicted fitness regions. Low temperatures, which decreases acceptance probabilities, produced overly exploitative trajectories that had monotonically increasing fitness traces. A temperature of 0.01 produced trajectories with fitness traces that on average improved but were not monotonic, suggesting a qualitatively good exploration-exploitation balance.

For the prospective GFP designs presented in the main text we used a trust radius of 15 mutations, and for a smaller scale experiment presented in Supplementary Figure 4, we used a trust radius of 7 mutations. For the prospective TEM-1 β-lactamase designs we used a trust radius of 7 mutations. We reduced the trust radius relative to GFP because only single mutants were used as low-N training data for the TEM-1 β-lactamase experiments.

From here, final sequence proposals were obtained by filtering the 3500 × 3000 = ∼10 million sequences explored for each independently trained sequence-to-function model. This was done by finding the best sequence in each trajectory and then selecting the top P sequences among these best-in-trajectory selections, where P=300 was the design budget. We did not do any further filtering to ensure mutual diversity as the selected sequences were already diverse in terms of pairwise number of mutations apart.

### Library Cloning and Transformation

For library cloning and transformation, we assume that we had available as input the output of a PCR reaction, where the 5’ and 3’ ends contain TIIS restriction sites compatible with golden gate assembly. For SynNeigh and FP Homologs, this corresponded to error-prone PCR product made with primers with appropriate TIIS flanking sequences. For each prospectively designed GFP and TEM-1 β-lactamase variant, corresponding DNA oligos contained 5’ and 3’ primer sequences such that their corresponding oligo pools could be amplified. Internal to these priming sequences were TIIS restriction sites that would cut internally into the oligo containing the coding sequence of the variant, and would consequently “clip off” the priming sequences.

All library clonings and transformations were performed using the following general steps: 1) PCR of the vector backbone, 2) golden gate assembly of the insert and vector, 3) ethanol precipitation of the ligated plasmid, 4) electroporation into electrocompetent DH5α *E. coli*, recovery, and subsequent outgrowth under selection.

Vector PCRs were performed with primers adjacent to the insert region that extended into the vector backbone. Vector primers were also adapted with TIIS restriction sites (either BsaI or BbsI) such that 4bp complementarity would be achieved with the library (“insert”) on both the 5’ and 3’ end after digestion with the appropriate TIIS enzyme. Vector PCRs were performed using Q5 High-Fidelity 2X Master mix (New England Biolabs). All GFP related libraries were cloned using BsaI sites. The prospectively designed TEM-1 β-lactamase library was cloned using BbsI sites. Both insert and vector PCRs were bead purified using homemade SPRI beads^80^.

PCRed vector and library inserts were then cloned using a one-pot Golden Gate Assembly reaction that contained TIIS restriction enzyme (BsaI-HF-v2 or BbsI-HF), T4 DNA ligase, and DpnI. Reactions were cycled between 37 **°**C and 23 **°**C to encourage iterative cutting and ligation. All enzymes were ordered from New England Biolabs. Reactions were then ethanol precipitated to purify the ligated plasmid in a form suitable for high-efficiency electroporation, and then electroporated into DH5α *E. coli* (Lucigen 10G Elite) cells using 0.1 cm electroporation cuvettes (GenePulser cuvettes, Bio-Rad) and a Bio-Rad MicroPulser. Electroporations were recovered in 1 mL recovery media (Lucigen) for 1 hour and subsequently grown overnight in LB + selection.

### FlowSeq

Our FlowSeq procedure was adapted from Kosuri *et al*. (2013)^81^. For every FlowSeq experiment we followed these steps:

Set up:

1. The night before, we grew up 1mL cultures of the following control strains: DH5α *E. coli*, DH5α *E. coli* expressing avGFP, and DH5α *E. coli* expressing sfGFP.
2. 500 uL of the library (either frozen stock or outgrown transformation from the night before) was diluted 1:100 into 50 mL of LB + selection, and shaken at 37C. Control strains were handled similarly at smaller scale.
3. Once cells for both the library and control strains reached OD_600_ of 0.1-0.4, cultures were washed 2x in 1X ice cold PBS buffer.
4. Control avGFP and sfGFP strains were “spiked” into the library at a representation of 0.1% to serve as internal standards.
5. Cells were passed through 100 micron cell strainer and were kept on ice for 2 hours. Fluorescence activated cell-sorting (FACS):
6. All FACS were performed on a Sony SH800S cell sorter. Unless otherwise noted, all excitation lasers (405 nm, 488 nm, 561 nm, 638 nm) were turned on, and readings were taken and gates were drawn with respect to filter FL2 (525/25 nm). Thus, only the 405 nm and 488 nm lasers were relevant. We note that the FL2 measurement represents the emission induced by joint excitation with the 405 nm and 488 nm lasers.
7. We first flowed DH5α *E. coli* to determine FSC and SSC sensor gains and trigger thresholds. Using additional information from area and height FSC and SSC measurements, we drew a polygon gate to capture ∼90% of singlet events, excluding likely doublets.
8. We next flowed the avGFP and sfGFP control strains to adjust the FL2 sensor gain such that there was good dynamic range between the non-fluorescent DH5α and the fluorescent avGFP and sfGFP, without saturating the upper detection range. We confirmed the avGFP and sfGFP showed about 1 log_10_ difference in relative fluorescence. Finally, we flowed the library to confirm that its range of fluorescence values was well captured under these sensor settings.
9. We next drew B perfectly adjacent but non-overlapping gates or “bins” to partition the entire range of fluorescence values observed across FL2 for the library. For generating the SynNeigh dataset B=17. For FPHomologs B=8, and for the prospectively designed GFP library (Figure 2 of main text) B=8. The uppermost bin was always set such that it captured the upper tail of the fluorescence distribution. Bin minimums and maximums were noted.
10. Library variants in each bin were then collected using two-way sorts. Sorts were done into polystyrene tubes filled with 1 mL of LB + selection media, and we noted the number of events that were sorted into each bin.
11. Sorted cells for each bin were then added to 10 mL of LB + selection media, and grown overnight. Unused library (input into the FACS) was pelleted and frozen at -20 **°**C Next generation sequencing (NGS):
12. Cultures of each bin as well as the input library (hereafter, “input”) were mini-prepped (Qiagen).
13. Illumina sequencing ready amplicons of the library region (SynNeigh and prospectively designed GFP library) or barcode region (FP Homologs) of each sample were prepared using a two stage PCR strategy. Sample multiplexing and pooling was accomplished with a standard dual indexing strategy.
14. The amplicon pool was then bead purified with homemade SPRI beads and quality controlled with TapeStation analysis and with qPCR to ensure the final pool was properly indexed, of the right length, and accurately quantified.
15. When generating the SynNeigh dataset we used a MiSeq 2 × 300 bp V3 run directly sequence the ∼500 bp library region of GFP. When generating the FP Homologs dataset we used a NextSeq 2 × 75 bp mid-output run to sequence variant barcodes. When sequencing the prospectively designed GFP library, we sequenced the ∼280 bp library region using a NextSeq 2 × 150 bp mid-output run. Data-processing and log_10_(relative fluorescence) inference:
16. After sample demultiplexing, if multiple lanes were used during sequencing (NextSeq runs), their corresponding fastq files were pooled.
17. For each sample, read pairs were merged using FLASH v1.2.11^82^.
18. For each merged read in each sample, the library region or variant barcode was extracted using a regular expression that identified delimiting constant primer sequences used for preparing the amplicon sequencing pools.
19. For each extracted region in each sample, protein sequences were determined by translating the directly sequenced or associated (in the case of variant barcodes as done for FP Homologs) nucleotide sequence.
20. For each sample, the count of every unique protein sequence was then determined. And the total collection of unique protein sequences across all samples was used to create a variants x bins count table, C.
21. Using the metadata collected during the FACS we could then infer the log_10_(relative fluorescence) values of each variant using the following procedure:
  a. Compute relative abundance table, R, by dividing the columns of C by their sums. The columns of R sum to 1.
  b. Divide each column of R element-wise by the input relative abundance vector (relative abundance of variants in the library before FACS) to obtain a fold change table, F.
  c. Divide each row of F by its sum to obtain a table of adjusted abundances, A. Each row of A sums to 1.
  d. Each row of A, which corresponds to data for a particular protein variant, defines a discrete probability mass function over which FACS bins the variant will appear. We therefore set the he inferred log_10_(relative fluorescence) of variant i to be the median of the distribution A_i_.

### Ancestral Sequence Reconstruction

We used the FastML web server to perform ancestral sequence reconstruction (ASR)^43^. A version or release was not available, but the tool was used on October 21, 2019. As input, we provided a multiple sequence alignment of *Aequorean* FPs. Default FastML parameters were used otherwise: Phylogenetic tree reconstruction method = RAxML, Model of substitution = JTT, Use Gamma Distribution = Yes, Probability cutoff to prefer ancestral indel over character = 0.5.

Through examining the reconstructed phylogenetic tree, we isolated two interesting ancestral nodes N1 and N11. N1 was the ancestor for all sequences, whereas N11 was an ancestor that excluded the *Aequorea macrodactyla* sequences TagCFP, OFPxm, and TagGFP, which contain a large number of mutations relative to avGFP. From each node, we generated the top 5 most likely ancestral sequences at both N1 and N11. Because we were comparing ASR to model-guided approaches, ASR mutations outside of the 81 amino acid library regions were converted back to wild-type. These designs were submitted as a Gene Fragments order to Twist Biosciences and cloned individually with Gibson assembly (reagents from New England Biolabs).

### Consensus Sequence Designs

Consensus sequence design attempts to sample the most probable sequences given a position weight matrix (PWM). We generated a PWM using the same sequence alignment we used for ancestral sequence reconstruction. To sample the highest probability sequences from the PWM we used a Metropolis-Hastings sampler to explore 180,000 sequences from which we filtered the top 5 highest probability sequences. Repeated runs of this procedure as well as multiple rarefaction analyses showed that we consistently captured the top two most probable sequences (manually derived) and that beyond 180,000 explored sequences no further improvements in sequence probabilities would be observed. The top 5 consensus sequence designs were submitted as a Gene Fragments order to Twist Biosciences and cloned individually with Gibson assembly (reagents from New England Biolabs).

### Fitness determination for TEM-1 β-lactamase variants

For each concentration of ampicillin (0, 250, 1000, 2500 μg/mL) and for each biological replicate, we prepared 3 large 150 mm plates of LB agar + ampicillin. We then prepared overnight starter cultures of two biological replicates of the cloned designed library and wild-type TEM-1 β-lactamase. On the day of the experiment, we back-diluted starter cultures 1:100 and let them grow to OD_600_=0.5 at which point we placed them on ice. Cells were then washed 2x in ice cold 1X PBS, and the wild-type strain was spiked into the library cultures at 0.1%. 250 uL (about 600M) cells were spread onto each prepared plate. Plates were incubated at 37 **°**C overnight.

The next day plates were “scraped” by adding 1 mL of 1X PBS and 5-10 cell spreader beads. Plates were shaken laterally so beads could dislodge colonies and mix cells into the PBS. This cell mixture was pooled for the three replicate plates for each antibiotic condition and biological replicate. These were then pelleted, mini-prepped, and NGS sequenced in the same way as done for FlowSeq. A 2 × 150 bp NextSeq run was used to sequence the library region. A design’s fitness at a particular strength of antibiotic selection was determined to be the ratio of its relative abundance under selection to its relative abundance under no selection.

### Qualitative inference of k_cat_ and K M^-1^ changes for TEM-1 β-lactamase variants

At 250 μg/mL, we didn’t observe a difference in growth rate in cells expressing wild-type TEM-1 β-lactamase in liquid culture. At 2500 μg/mL we saw strong inhibition. Consequently, we assume these represent low ([S] < K_M_) and high ([S] > K_M_) substrate concentrations, respectively. We also assume that growth rate is proportional to the reaction velocity of ampicillin hydrolysis by the TEM-1 β-lactamase enzyme. From previous work, we know this hydrolysis reaction is well modeled by Michaelis-Menten dynamics^83^. The Michaelis-Menten equation is given by, reaction_velocity = k_cat_[E][S]/(K_M_ + [S]).

At high substrate concentrations, the reaction velocity is approximated by the expression k_cat_[E]. Variants with higher fitness in the high-substrate, 2500 μg/mL condition have higher abundance (controlling for their input abundance), which must be the result of a faster growth rate. Assuming that mutations we make to the enzyme do not change its expression and concentration inside the cell, [E], this in turn implies that these variants have an increased k_cat_. Cluster 1 designs exhibited this behavior (Fig. 3c). It straightforwardly follows that variants with lower fitness at 2500 μg/mL ampicillin have a lower k_cat_ (Cluster 2-4 variants, Fig. 3c).

At lower substrate concentrations, the reaction velocity is approximated by the expression k_cat_[E][S]/K_M_. Taking the ratio of this expression for a mutant enzyme and a wild-type enzyme we have, [k_cat_(mut) / k_cat_(WT)] x [K_M_(WT) / K_M_(mut)]. When a variant has higher fitness at the low-substrate 250 μg/mL condition, this ratio is greater than 1. Now if from the high-substrate condition we could infer that k_cat_(mut) < k_cat_(WT), then it must be the case that K_M_(mut) < K_M_(WT). This logic applies to Cluster 2-4 designs in Figure 3c. However, if from the high-substrate condition we inferred that k_cat_(mut) > k_cat_(WT), then without further information, we cannot guess the direction of change for K_M_(mut), which is the case for Cluster 1 designs.

### Exploration of evolutionary, structural, and principal component mutational patterns in designs

In our examination of the mutational patterns in proposed and successful designs we began by gathering high-quality Position-Specific Scoring Matrices from the ProteinNet database^84^ for both avGFP (PDB: 2WUR) and TEM-1 β-lactamase structure (PDB: 1ZG4). These PSSMs are without gaps. We computed the “effective number of mutations” per residue within our design window by taking the exponent of the per-position Shannon entropy, e.g. 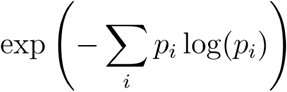. For residues where only one amino acid was observed in the multiple sequence alignment, the PSSM had 1 in that amino acid’s position and zero elsewhere, such that the effective number of mutations was 1. Likewise, if all amino acids were observed with equal frequency at that position, the effective number of mutations was 20.

For each position in the design window, we computed the relative frequency of mutation for the proposed and functional eUniRep designs. We counted the number of times a position was mutated to any residue outside the wild-type, and divided it by the total number of mutations for each set.

We computed a least-squares regression between the mutation tolerance and relative mutation frequency using Scipy (https://docs.scipy.org/) including the r-value and *p*-value (Fig. 4a-b, left). We also visualized the scatter plot of relative mutation frequency in proposed and gain of function designs along with the effective number of mutations (Fig. 4a-b, right).

Next we used the experimentally determined crystal structures for both proteins to analyze relationships between mutation frequency and structural features. We first examined the euclidean distance in 3-D space between the positions in the design window of avGFP and the centroid of the chromophore of avGFP (S65, Y66, G67). Likewise, we computed distances of positions within the design window of TEM-1 β-lactamase with the catalytic Serine S70’s side chain oxygen. Instead of examining the per-position distance, we took all bright designs and computed the distribution of distances of all the mutated position within each design, and visualized the relationship between the quantitative function score (log_10_(relative fluorescence) and log_10_(fitness)) and the mean distance of mutated residues from the active site along with 5th and 95^th^ percentile distances, computing a least squares regression, r-value, and p-value as above.

Using DSSP^85^, we inferred per-position secondary structure annotations and relative solvent accessibility. For the small residues without a DSSP annotation, we manually examined the crystal structure and classified the residues secondary structure by eye. All positions with relative solvent accessibility less than 0.2 were classified as buried, and all others were exposed^86^. We visualized the frequency of mutations in our design window into each secondary structure category if we were to mutate uniformly randomly, the null expectation, and compared it to the mutation frequency we observed in proposed and >WT eUniRep designs (Fig 4c-d, bottom). We colored the crystal structures of each protein by the relative per-position mutation frequency in >WT designs (Fig 4c-d, upper center).

Lastly, we examined the relationship between function and the euclidean space defined by eUniRep’s vector representation. We sampled sequences with a random number of mutations ∼ Poisson(4) + 1 (uniform across the sequence length) relative to wild-type for both proteins. eUniRep representations were computed for each, along with one-hot encoded matrices. We performed principal component analysis on the representations of this collection of random sequences, and subsequently projected representations of the experimentally characterized random mutant sequences of avGFP from Sarkisyan *et al*. (2016)^42^ and the single mutants of TEM-1 β-lactamase from Firnberg *et al*. (2014)^48^ onto the first and second PCs of both eUniRep (avGFP and TEM-1 β-lactamase) and Full AA (avGFP and TEM-1 β-lactamase). Projected sequences points were colored by their quantitative function. We computed Pearson’s correlation between the measured quantitative function and eUniRep PC1, as well as Full AA PC1.

Each model’s ability to differentiate top >WT designed sequences from WT on the basis of predicted function (Fig. 4h-i), was defined to be the (signed) number of standard deviations predicted wild-type function was from the median of the top sequence design predicted functions. For robustness, standard deviation was estimated using the median absolute deviation.

