## Supplementary Information for "Low-N protein engineering with data-efficient deep learning"

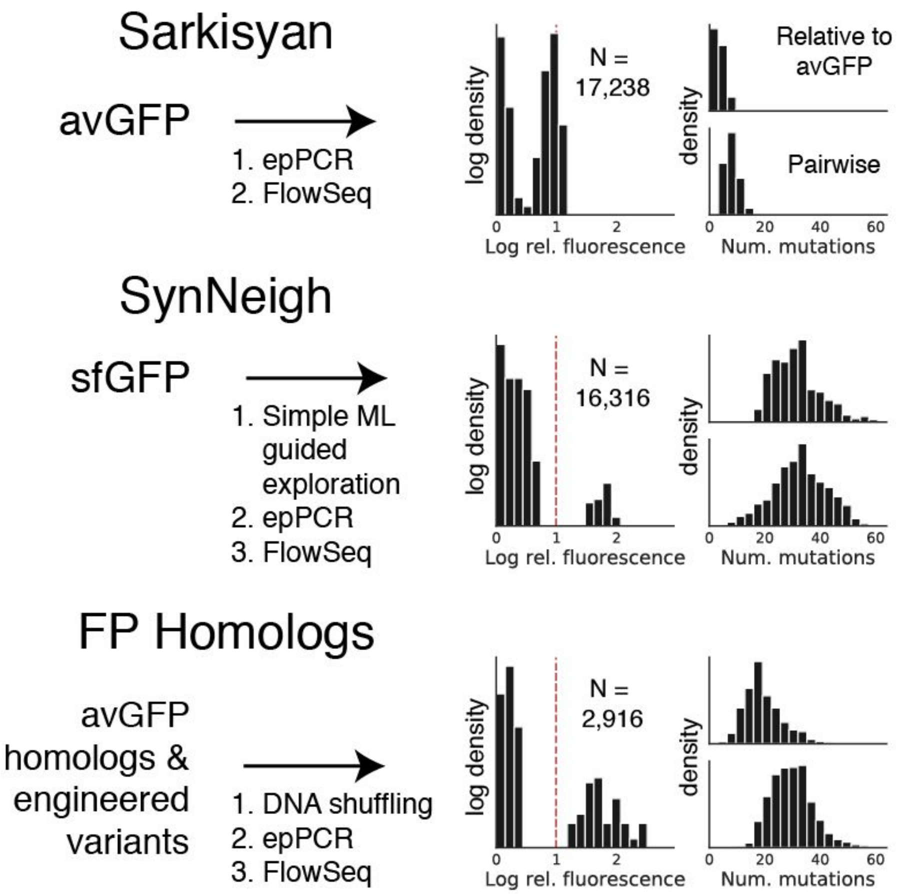


**Supplementary Figure 1.** Summary of datasets used for retrospective experiments. Detailed descriptions can be found in the Methods. For each dataset -- Sarkisyan, SynNeigh, and FP Homologs -- shown are the starting sequence(s) and how it was manipulated to obtain the final dataset. Plots on the right illustrate summary statistics that include the distribution of relative fluorescence values, the distribution of the number of mutations each variant carries with respect to avGFP, and the distribution of pairwise mutation distances for the dataset. Sarkisyan was processed from Sarkisyan *et al.* (2016)[42] and served as a source of low-N training sets. SynNeigh and FP Homologs were generated and processed in this work and served as generalization sets to ascertain low-N generalization performance.

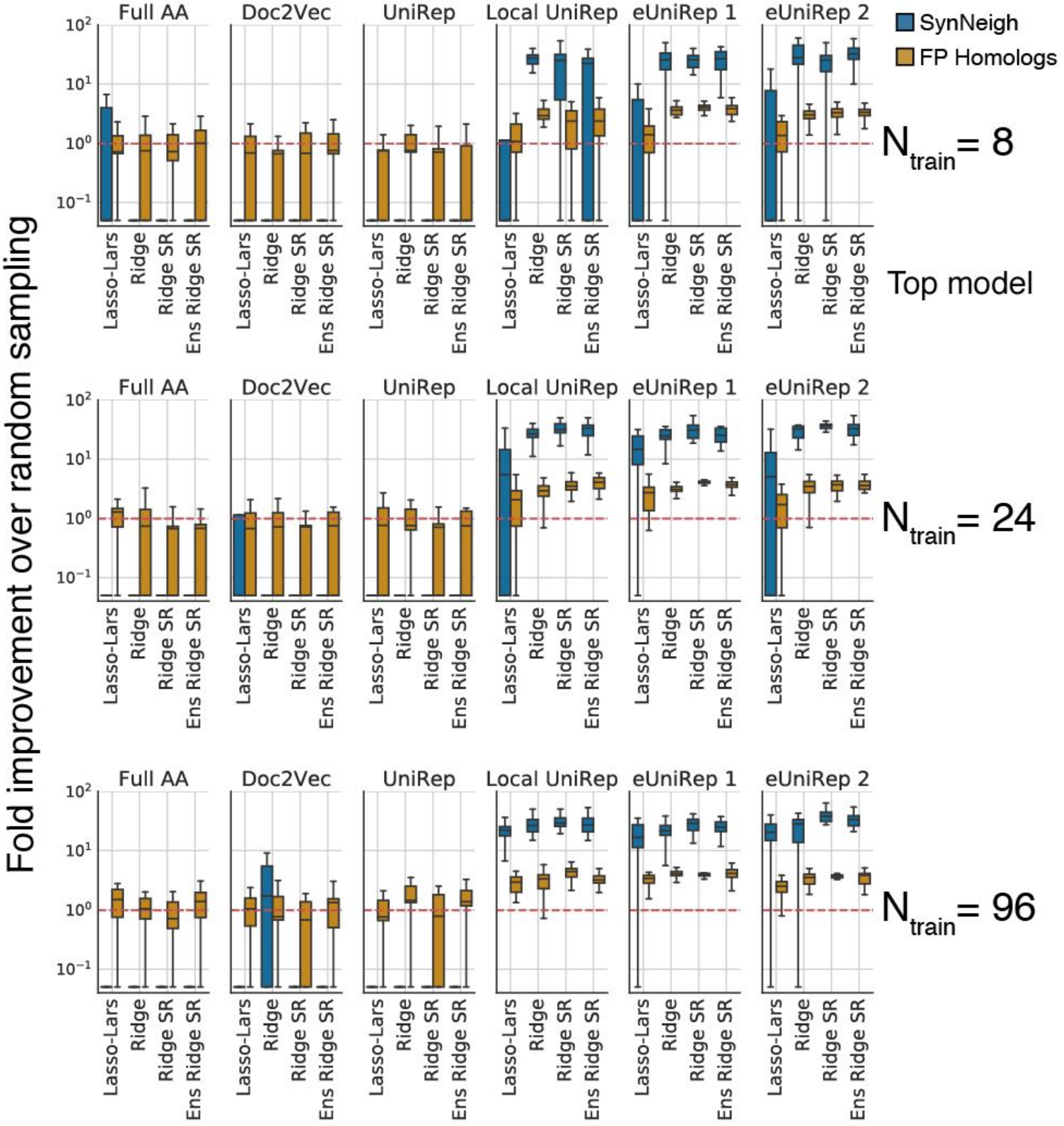


**Supplementary Figure 2.** Summary of retrospective performance for different numbers of training sequences, choices of sequence representation, and choices of top model. A complete description of the experiments performed can be found in the Methods. Briefly, given a low-N trained sequence-to-function model (training data from Sarkisyan) we evaluate its generalizability on held out sequences in a generalization set (SynNeigh or FP Homologs). Generalization is measured by the ability of this model to rank order sequences in the generalization set such that when the top 96 are selected, as many as possible are >WT. The y-axis therefore measures the number of >WT variants found among the top 96 ranked divided by the average of the same with respect to 1000 random orderings. Results reported on Split 1 of retrospective datasets (Methods).

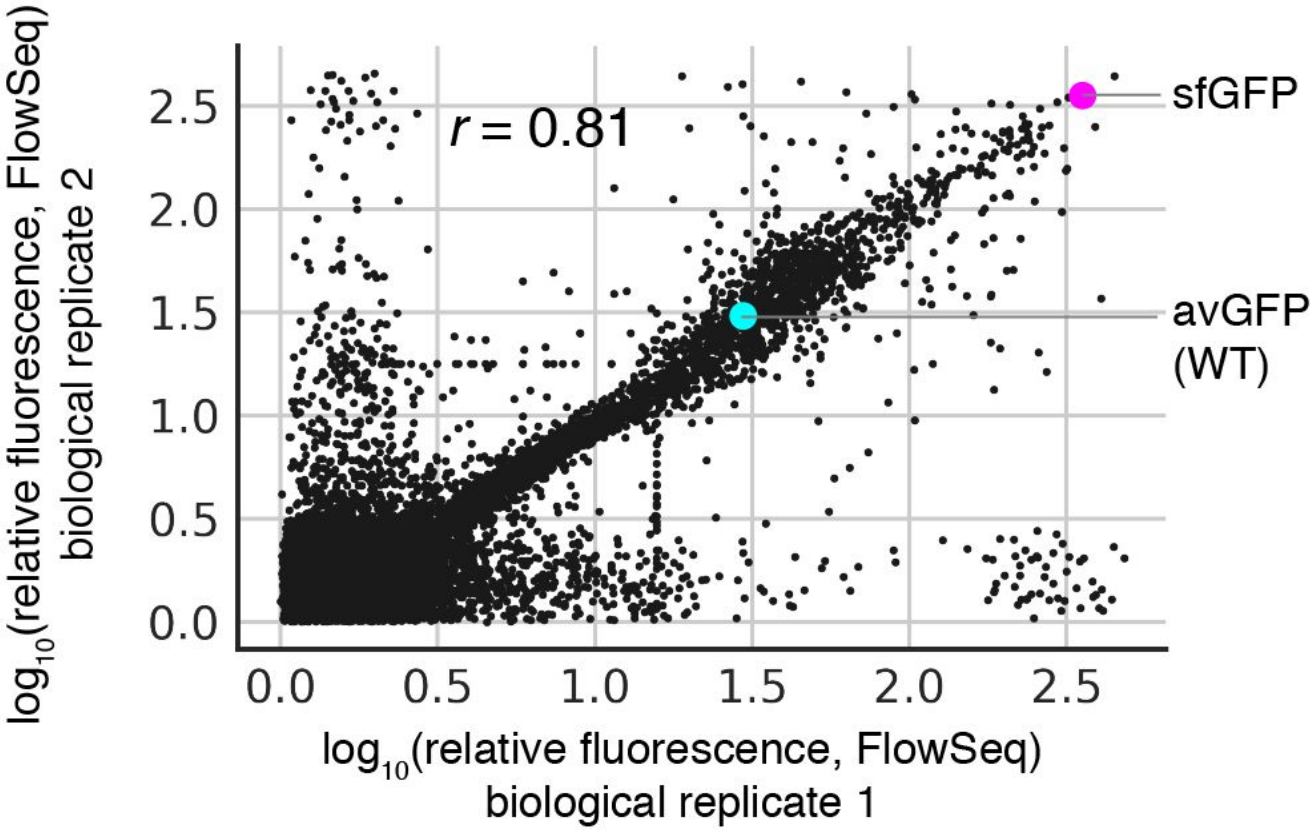


**Supplementary Figure 3.** Biological reproducibility of FlowSeq assay for all prospectively designed GFP variants shown in Figure 2. Each biological replicate consisted of independent cloning, transformation, and FlowSeq steps. Pearson correlation between the two replicates was 0.81. avGFP (cyan) and sfGFP (magenta) are shown. Replicates were conservatively pooled by taking the minimum value of replicate measurements for each designed variant.

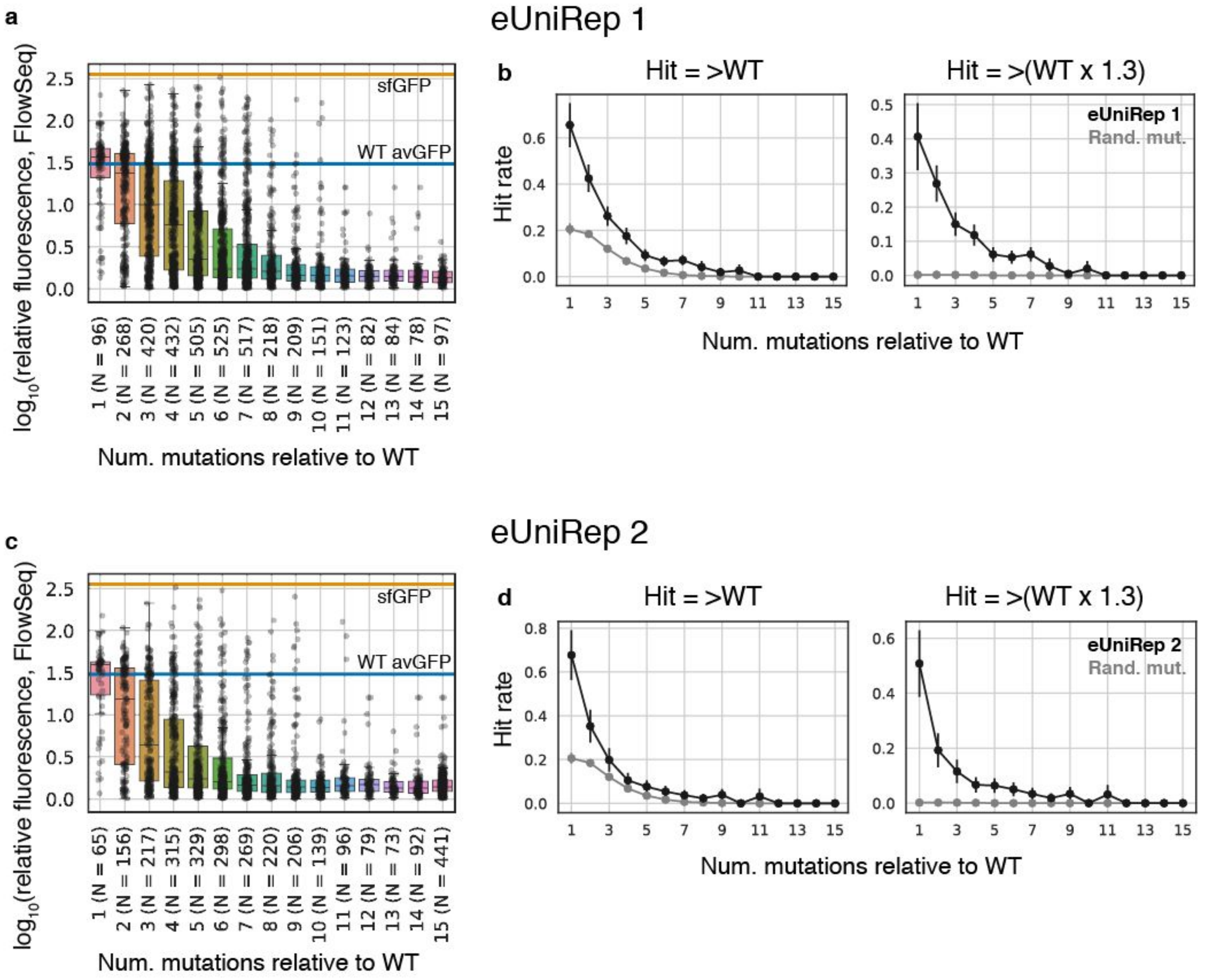


**Supplementary Figure 4.** Protein function as measured by fluorescence and hit rate as a function of number of mutations relative to wild-type avGFP. **a)** $\log_{10}$(relative fluorescence) of variants as a function of the number of mutations they have relative to wild-type avGFP. Also noted are the number of designs containing each number of mutations. **b)** Hit rates as a function of number of mutations relative to wild-type where a "hit" is defined either as a variant with >WT activity (left) or with greater than 1.3x WT activity (right). For context, hit rates for random mutagenesis are also shown. These were calculated from the error-prone PCR avGFP mutants measured in Sarkisyan *et. al*. (2016).

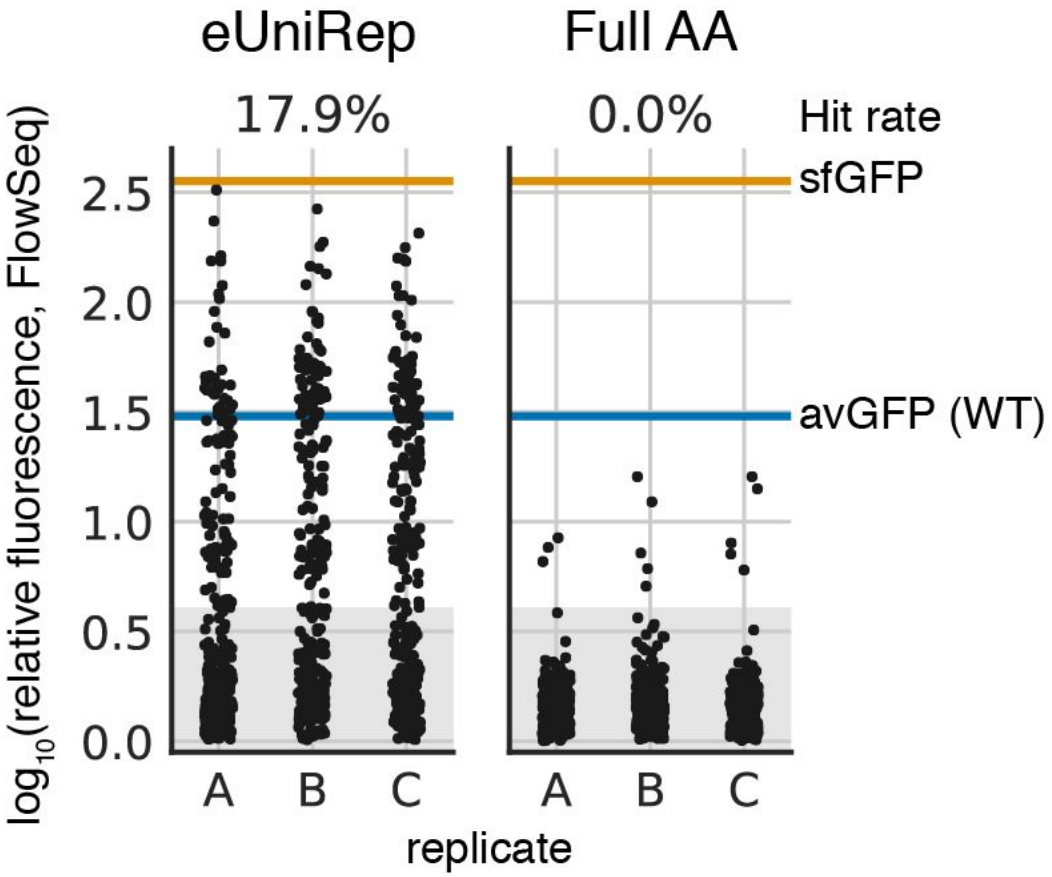


**Supplementary Figure 5.** Fluorescence intensities for prospective GFP designs when a smaller trust radius of 7 is used instead of 15.

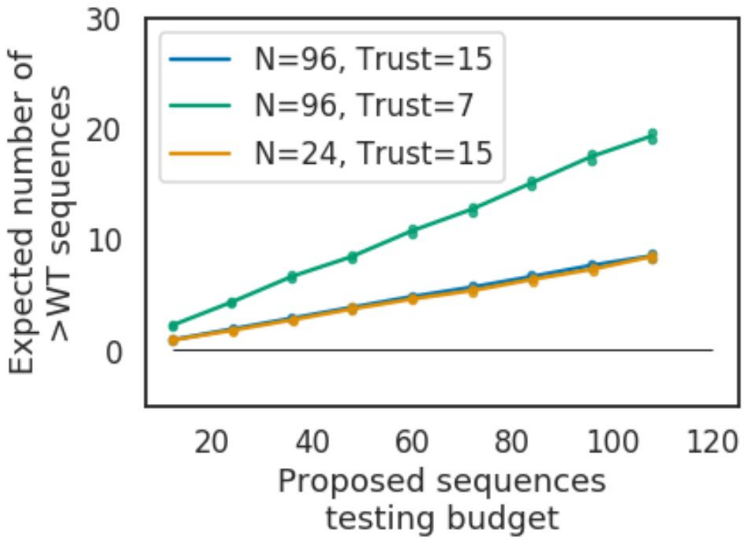


**Supplementary Figure 6.** Simulated performance of UniRep-based in silico directed evolution at lower testing budgets (12, 24,...120 testing points) for proposed GFP sequences. Means plotted as lines and 95%

Confidence Interval (CI) above and below as dots (shifted horizontally for visibility). At N=24 training points, 24 testing points appears sufficient to obtain at least one > WT variant (1.80 +/- 0.8 95% CI from 1000 bootstrapping samples). For each bootstrapping sample we picked experimental replicate and model replicate (one of the two eUniRep models) randomly, and then used the success rate of that experiment as parameter p of the Bernoulli distribution to generate simulated experimental outcomes to fill a given testing budget. We then used resulting bootstrap samples to obtain mean and 95% CI for the number of >WT sequences at that sequence budget.

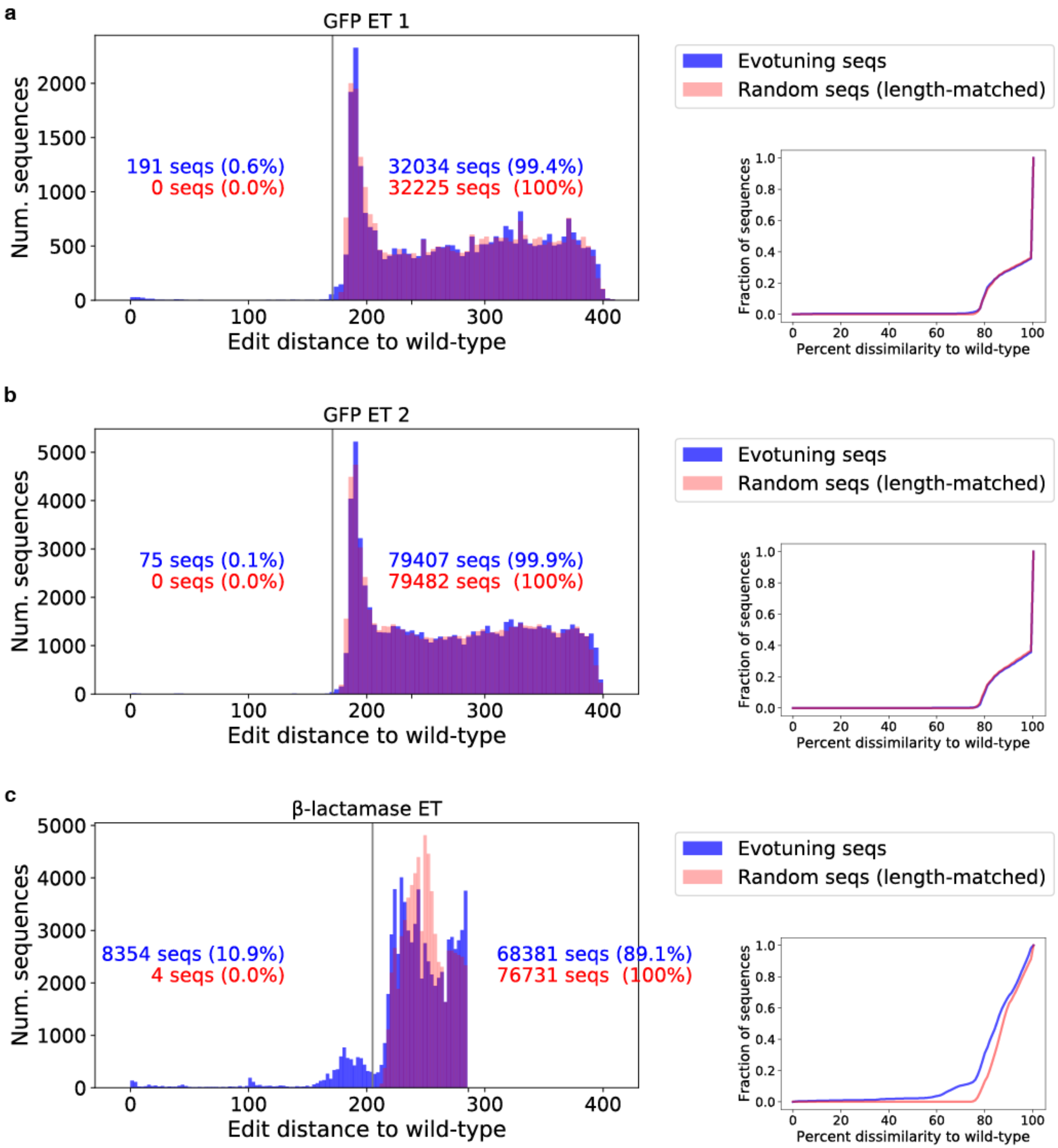


**Supplementary Figure 7.** Edit distance as computed by Levenshtein distance (equivalent to Needleman-Wunsch global sequence alignment with equally weighted penalties) of evotuning sequences (blue) to wild-type for **a)** GFP evotuing set 1, **b)** GFP evotuning set 2, and **c)** TEM-1 β-lactamase evotuning set. Vertical gray line corresponds to 72% sequence dissimilarity. Right inset plots illustrate the empirical cumulative distribution function of the sequence dissimilarity of these evotuning sequences to wild-type. Overlaid red histogram and line, respectively, depicts distribution of edit distances of random sequences from UniRef50 that are length-matched to those in the evotuning set.l The majority of sequences in all evotuning sets are indistinguishable from random, length-matched UniRef50 sequences on the basis of edit distance to wild-type.

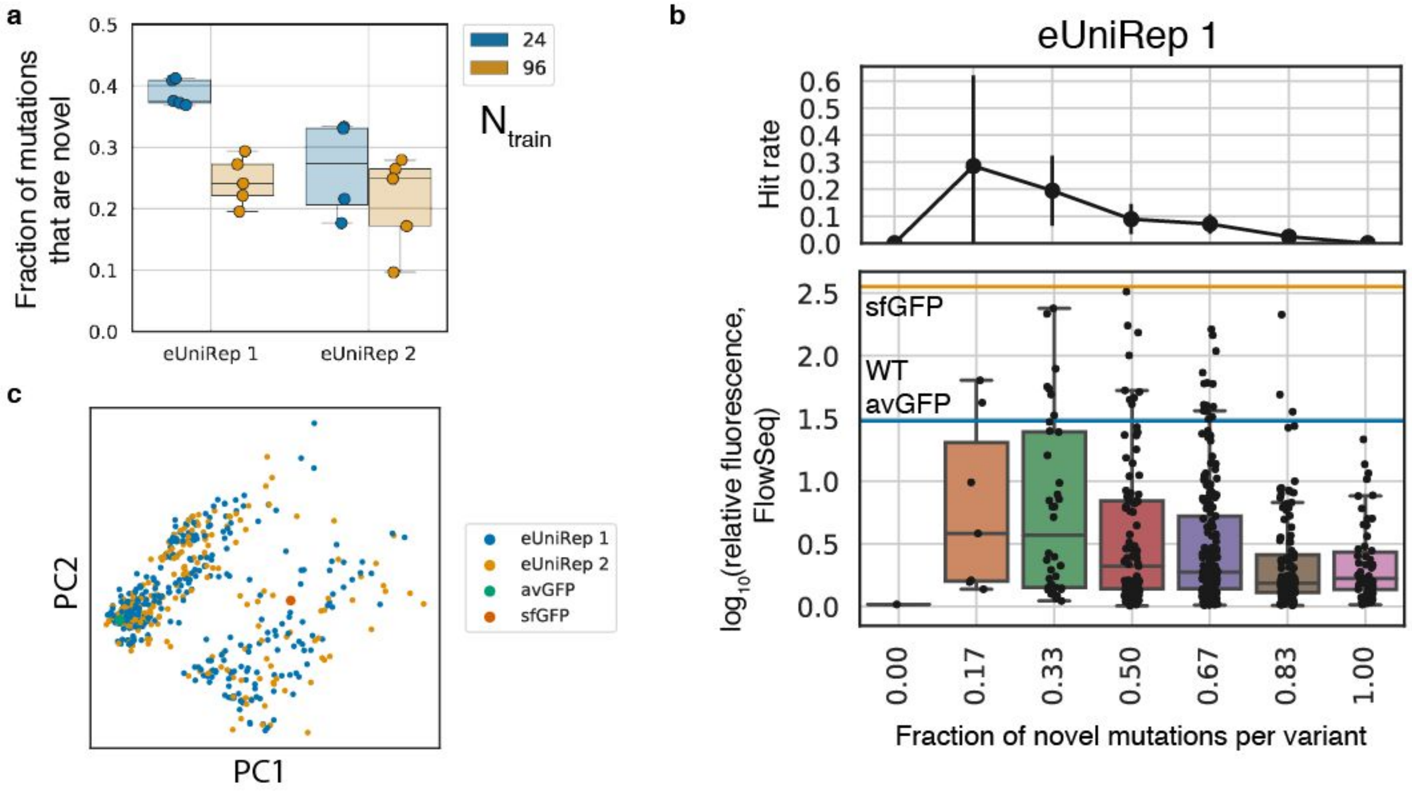


**Supplementary Figure 8.** Comparison of mutations found in >WT GFP sequence designs to those found in evotuning sequences and low-N training sequences. **a)** Fraction of mutations found in designed sequences that are not found in evotuning sequences or the low-N training sequences. Mutational overlap was tabulated after performing a multiple sequence alignment of each set of design sequences with their corresponding low-N training seqs and the appropriate evotuning set sequences. Each datapoint reflects a replicate's worth of designs, and replicates are partitioned by training set size (blue for 24, orange for 96). **b)** Hit rate and protein activity as a function of the fraction of novel mutations among all mutations each variant contains. Only variants (black dots) with 6 mutations are shown (the most common mutation count among all GFP designs). **c)** PCA plot illustrating similarity of >WT eUniRep 1 and 2 designs. Edit distances between sequences generated by a given model were similar to edit distances between sequences generated by different models (intra-eUniRep 1 distances = 6.3 +/- 2.5 s.d., intra-eUniRep 2 distances = 6.4 +/- 2.8 s.d., inter-eUniRep 1 & 2 distances = 6.4 +/- 2.6).

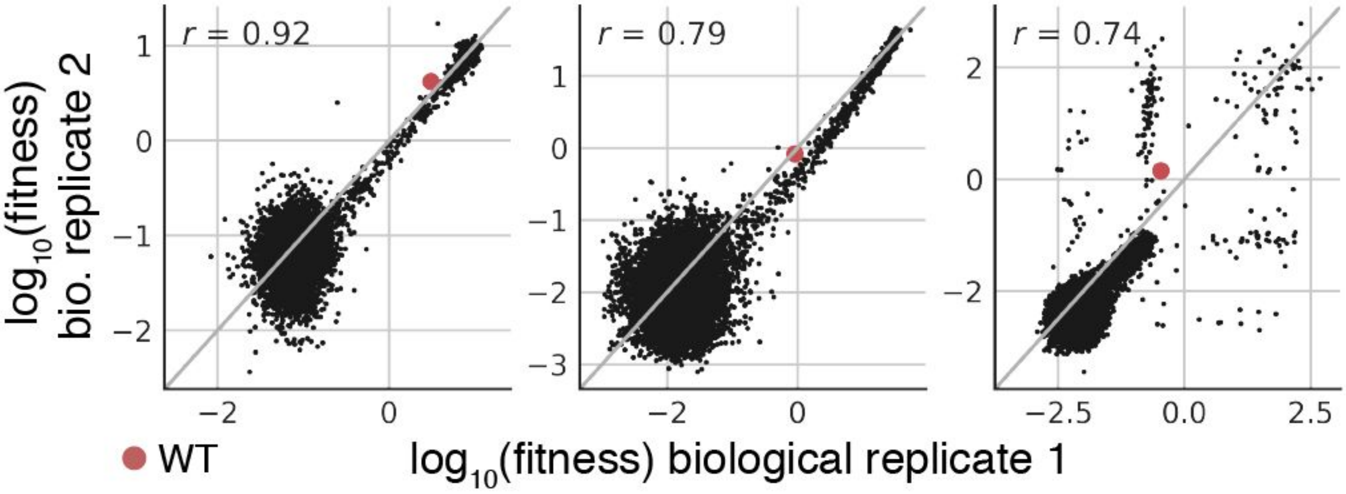


**Supplementary Figure 9.** Biological reproducibility of the fitness assay used for all prospectively designed TEM-1 β-lactamase variants shown in Figure 3. Each biological replicate consisted of independent cloning, transformation, plating, scraping, and NGS sequencing steps. Pearson correlation between the two replicates ranged between 0.74 and 0.92 depending on the strength of ampicillin selection. Wild-type (red circle) is shown. Replicates were conservatively pooled by taking the minimum value of replicate measurements for each designed variant.

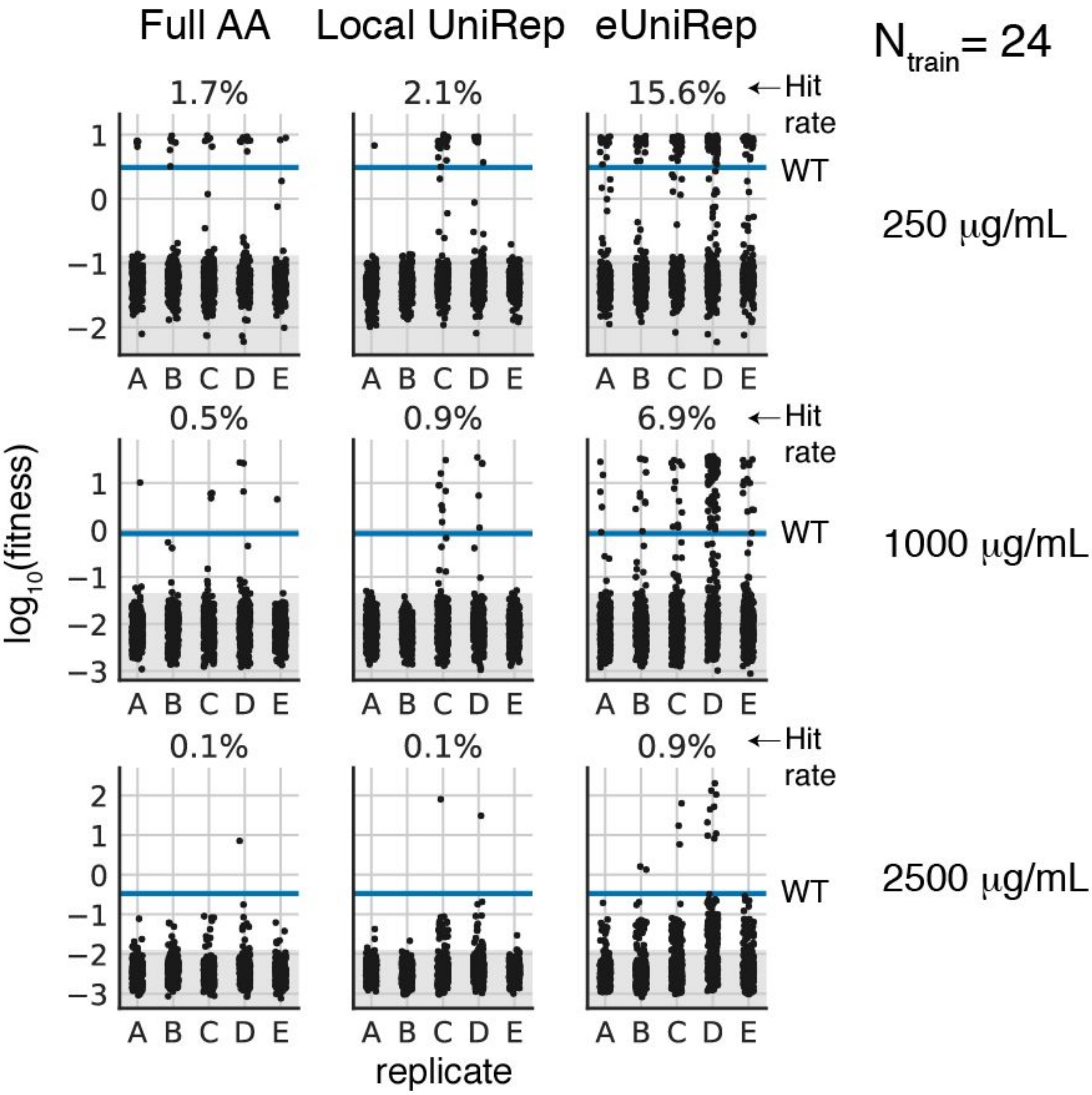


**Supplementary Figure 10.** Fitness of prospective TEM-1 β-lactamase designs when using training sets of size N=24, instead of N=96 as shown in Figure 3b.

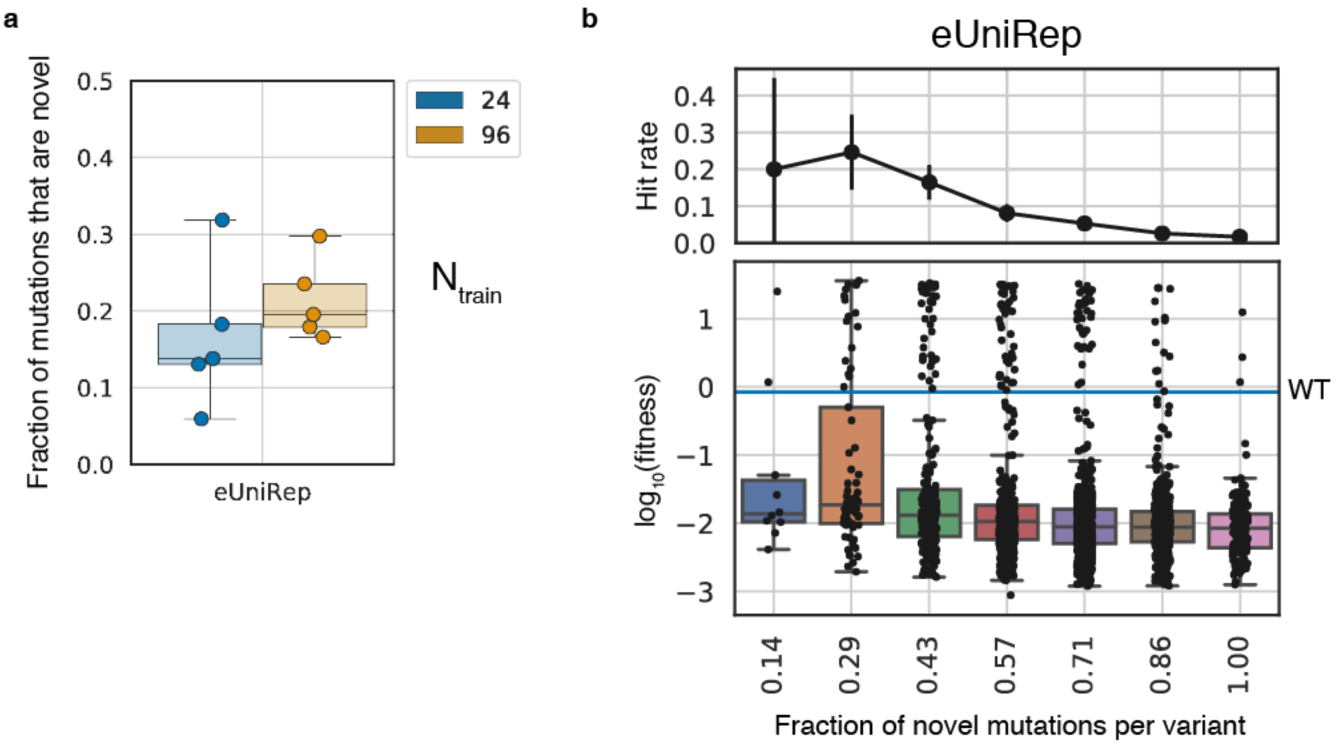


**Supplementary Figure 11.** Comparison of mutations found in >WT TEM-1 β-lactamase sequence designs to those found in evotuning sequences and low-N training sequences. **a)** Fraction of mutations found in designed sequences that are not found in evotuning sequences or the low-N training sequences. Mutational overlap was tabulated after performing a multiple sequence alignment of each set of design sequences with their corresponding low-N training seqs and the appropriate evotuning set sequences. Each datapoint reflects a replicate's worth of designs, and replicates are partitioned by training set size (blue for 24, orange for 96). **b)** Hit rate and protein activity as a function of the fraction of novel mutations among all mutations each variant contains. Only variants (black dots) with 7 mutations are shown (the most common mutation count among all TEM-1 β-lactamase designs).

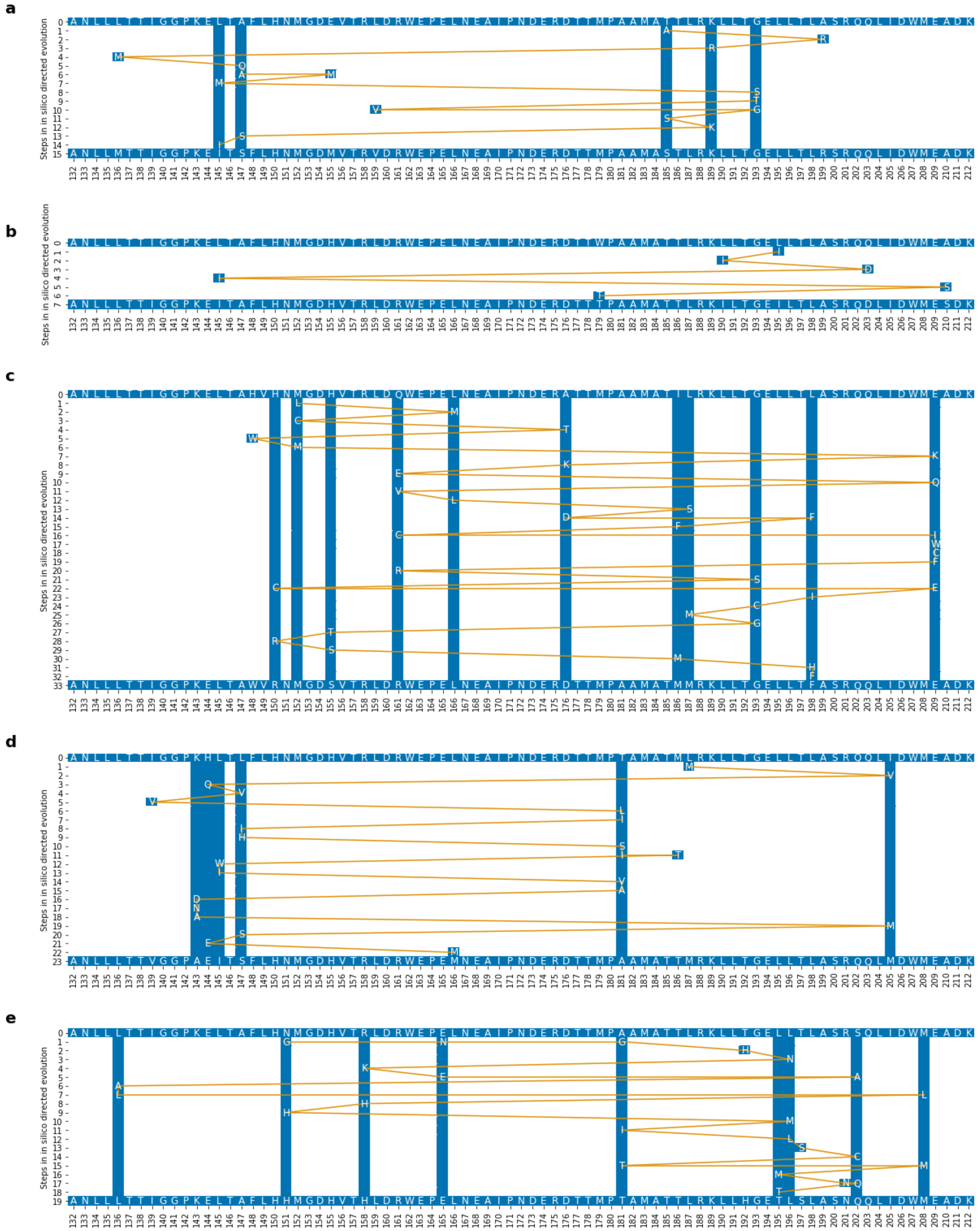


**Supplementary Figure 12.** Mutational trajectories in *in silico* directed evolution of 5 highly-functional TEM-1 variants predicted to be the most deleterious under additivity of single mutant effects. **a-e)** Top row in each panel represents the starting point TEM-1 variants were then optimized by the *in silico* evolution (81-amino acid region engineering region shown). Bottom row presents the same region of the terminal sequence. Yellow line demonstrates the accepted mutations along the trajectory from start to terminal sequence, with blue columns highlighting positions that were mutated more than once on the course of the trajectory. While some trajectories show straightforward accumulation of mutations (**b**), others display nuanced mutation ordering patterns (**a, c, d, e**). Sometimes, position X assumes a new amino acid, then other non-X positions are mutated, after which X amino acid is reverted to its initial amino acid (e.g. in **c**: initial M152 mutates to L152, then mutations occur in positions 166, 148, 176, after which 152 reverts back to M). This raises the possibility that UniRep-based *in silico* evolution navigates epistatic interactions that make some beneficial mutations inaccessible in certain sequence contexts.

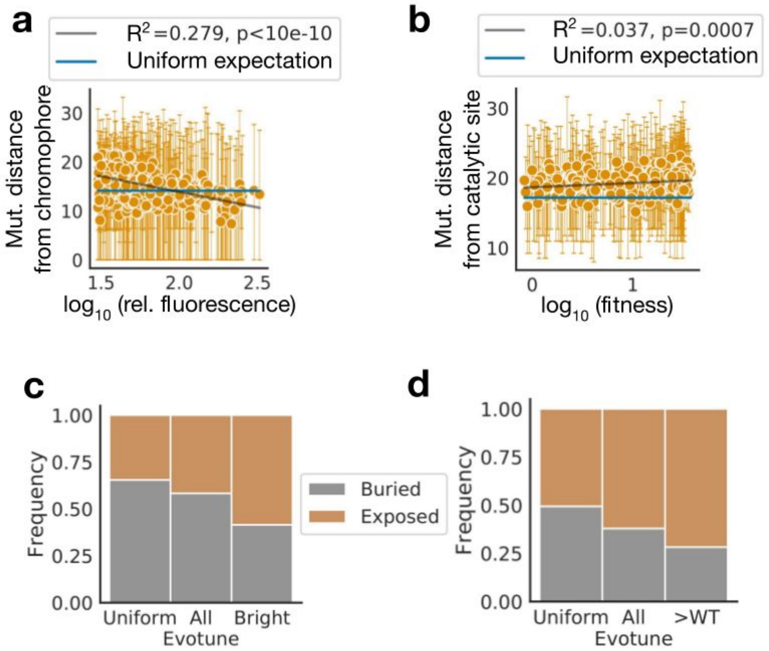


**Supplementary Figure 13.** Structural mutation patterns. **a)** Linear relationship between mean distance (Å) of mutations from chromophore, per variant. Dots are means and error bars 5th to 95th percentile. Blue shows expected distance sampling uniformly in the design window. **b)** Same as e) but for TEM-1 β-lactamase. **c)** Relative mutation frequency in buried vs. exposed residues (Methods) for avGFP, comparing a uniform expectation, all evotuned UniRep (eUniRep) designs, and the eUniRep designs which were greater than WT activity. **d)** Relative mutation frequency in buried vs. exposed residues, as in **c** (Methods) for TEM-1 β-lactamase.

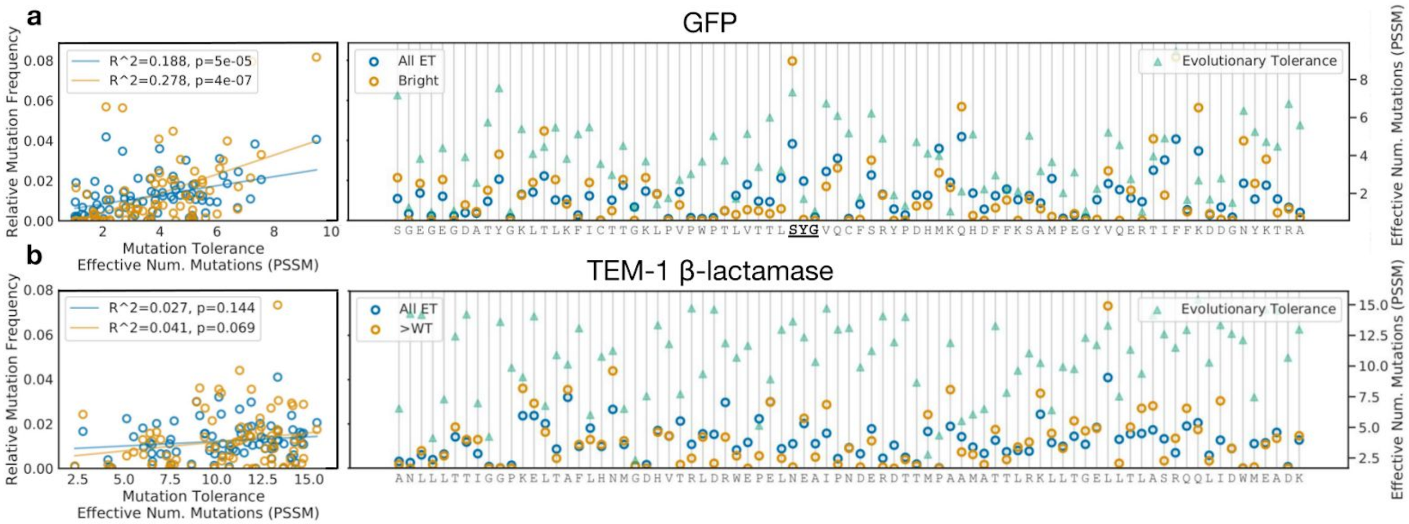


**Supplementary Figure 14.** Evolutionary tolerance and mutation frequency. **a)** (left) Relative mutation frequency per position as a function of mutational tolerance in N effective mutations (Methods). Blue are all mutations proposed across eUniRep designs. Orange are the subset of those mutations associated with >WT designs. (right) relative mutation frequency and effective N mutations over design window primary sequence. Chromophore bold and underlined. **b.** As in a but for TEM-1 β-lactamase, using $\log_{10}$(fitness) in the 1000 μg/ml condition for functional score.

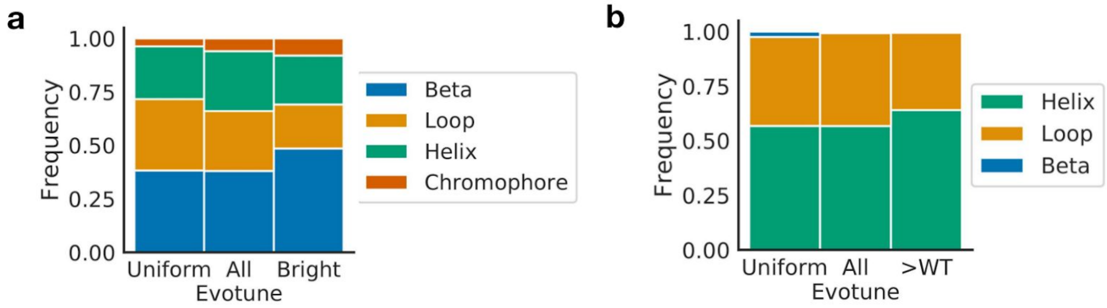


**Supplementary Figure 15.** Relative mutation frequency in secondary structure features for avGFP **a)** and TEM-1 β-lactamase **b)** Uniform shows expectation from uniform random sampling in design window, All Evotune are all eUniRep designs, and >WT are designs which are better than WT activity.

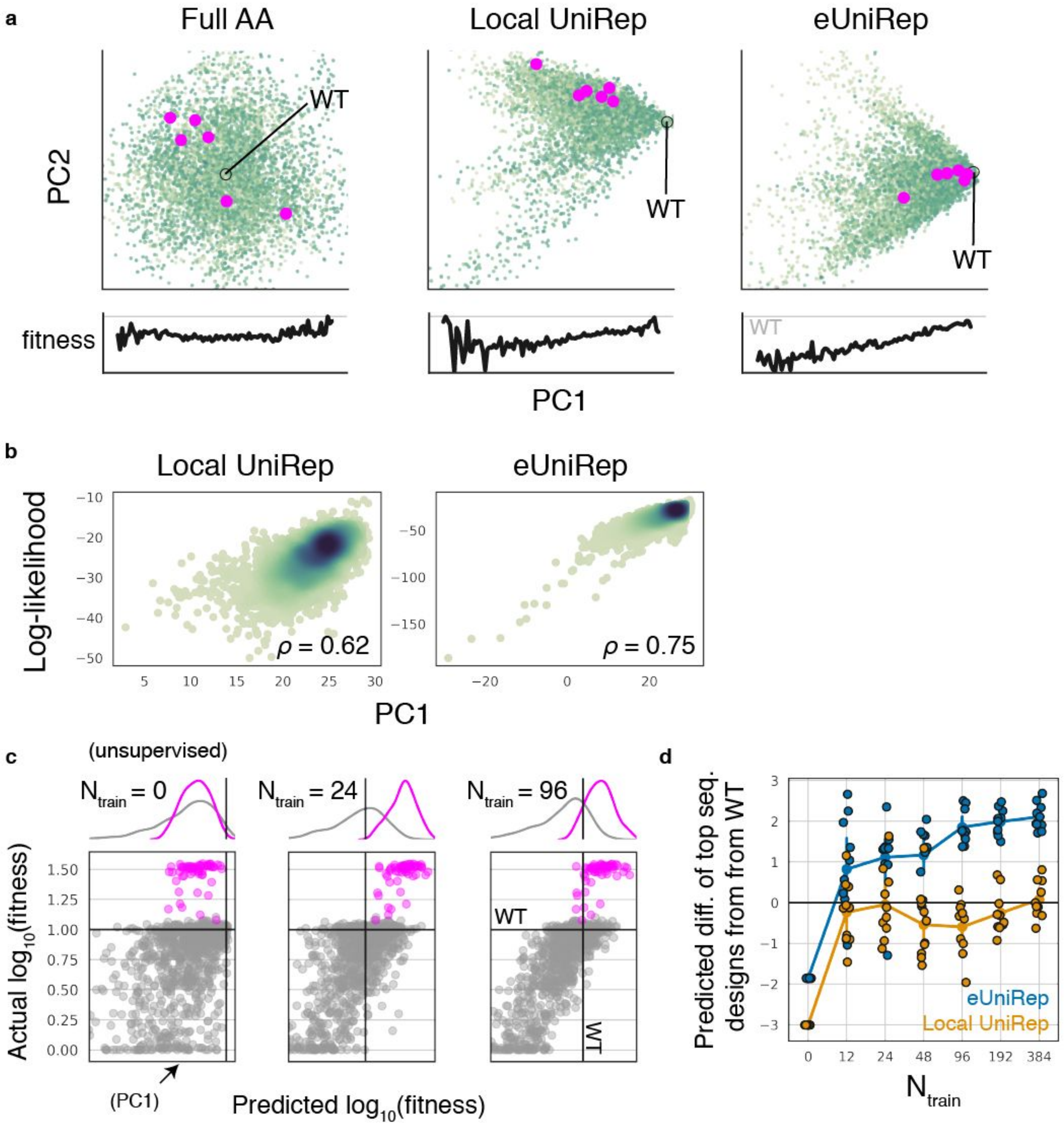


**Supplementary Figure 16. a)** PCA of Full AA, Local UniRep, and eUniRep representations of sequences from the local fitness landscape of TEM-1 β-lactamase colored by fitness. Note that this includes only single mutants. Magenta points show the top 10 sequence designs produced by each model. Wild-type TEM-1 β-lactamase is also marked. Below each plot, fitness as a function of PC1, Pearson $r = 0.05$ (Full AA), $r = 0.20$ (Local UniRep), $r = 0.44$ (eUniRep). **b)** Scatter plot of sequence log-likelihood under a given model (left Local

UniRep, right eUniRep) versus PC1 of its sequence representation. **c)** Scatter plots of actual vs predicted $\log_{10}$(fitness). $N_{train}$ = 0 corresponds to a purely unsupervised case, and so the x-axis is PC1. Grey circles are examples from the training distribution from which low-N training mutants are sampled. Magenta points represent the top 59 designed TEM-1 β-lactamase sequences. Kernel density estimates of each population are shown above each scatter plot. **d)** Jitter plot depicting the degree to which top sequence designs can be differentiated from wild-type on the basis of predicted activity as a function of the number of low-N training mutants used (Methods). At a given $N_{train}$, each datapoint represents a prediction replicate, which involves an independently sampled low-N training sequence set.

**a**

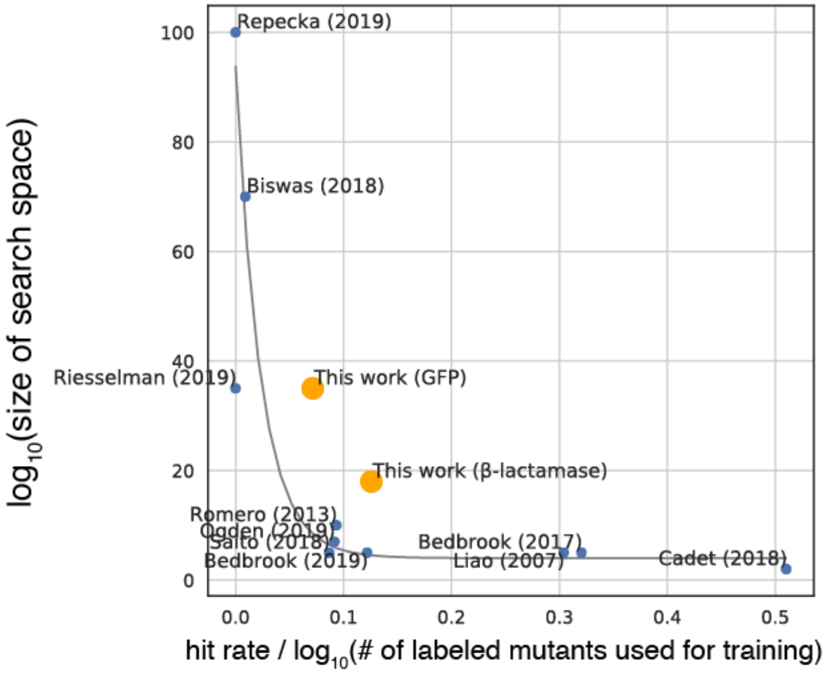


**b**

| Study | Search in sequence space | Requires structure? | Key achievement |
|---|---|---|---|
| This work (GFP) | extrapolative | no | -- |
| This work (β-lac.) | extrapolative | no | -- |
| Ogden (2019) | extrapolative | no | ML guided design of novel AAVs for gene therapy |
| Bedbrook (2019) | interpolative | yes | ML guided design of channelrhodopsins for minimally invasive optogenetics |
| Wu (2019) | interpolative | no | Enumeration of strategies for ML guided navigation of combinatorial libraries |
| Cadet (2018) | interpolative | no | Novel Innov'SAR approach for function prediction from sequence |
| Saito (2018) | interpolative | no | ML guided design of yellow-shifted FPs |
| Biswas (2018) | extrapolative | no | Suggests ML can be used to explore sequence space for directed evolution |
| Romero (2013) | interpolative | yes | First to use Gaussian processes for protein design |
| Bedbrook (2017) | interpolative | yes | ML guided design of channelrhodopsins that express and membrane localize |
| Liao (2007) | interpolative | no | Efficient ML guided design of the stability of proteinase K |
| Repecka (2019) | interpolative | no | First work to generate a functional protein with a GAN |
| Riesselman (2019) | interpolative | no | First work to generate nanobody sequences using a dilated causal conv net |

**Supplementary Figure 17.** eUniRep-guided low-N design demonstrates the strongest generalization performance shown to date. In this analysis, we define generalization performance to be how often a method discovers >WT sequences within a search region of a given size, while controlling for the number of training data mutants. We tabulated these quantities from this work and 11 previous machine learning guided-protein design studies. **a)** Scatter plot illustrating the relationship between the size of the search region each study explored and the study's hit rate normalized to the $\log_{10}$ number of functionally characterized mutants used for training. Hit rate is normalized to the size of the training set in this manner because we *a priori* expect hit rate would improve with larger training sets. Most studies were organized along a "front" (gray line), suggesting an inherent trade-off between the size of the search space and the likelihood of finding high-functioning variants (controlling for training set size). eUniRep-guided low-N design (this work) managed this trade-off substantially better achieving normalized hit-rates that other approaches require a $10^{15}$ to $10^{25}$ smaller search region for. **b)** Qualitative descriptors of the machine learning methods used in each study. Most studies were interpolative, such that their training sequence distribution overlapped with the sequence distribution they aimed to generalize to. The remaining studies, including ours, were extrapolative and generalized to regions of the fitness landscape beyond the training distribution. Additionally, a few of the approaches require structural data, whereas the others, including ours, do not. Finally, the key achievements of each study are included as reference.
